# Shield co-opts an RmuC domain to mediate phage defence across *Pseudomonas* species

**DOI:** 10.1101/2022.11.04.515146

**Authors:** Elliot Macdonald, Henrik Strahl, Tim R. Blower, Tracy Palmer, Giuseppina Mariano

## Abstract

Competitive bacteria-bacteriophage interactions have resulted in the evolution of a plethora of bacterial defense systems preventing phage propagation. In recent years, computational and bioinformatic approaches have underpinned the discovery of numerous novel bacterial defense systems. Anti-phage systems are frequently encoded together in genomic loci termed defense islands. Here we report the identification and characterisation of a novel anti-phage system, which we have termed Shield, that forms part of the *Pseudomonas* defensive arsenal. The Shield system comprises a membrane-bound protein, ShdA, harboring an RmuC domain. Heterologous production of ShdA alone is sufficient to mediate bacterial immunity against a panel of phages. We show that ShdA homologues can degrade phage DNA *in vitro* and, when expressed in a heterologous host, can alter the organisation of chromosomal DNA to a nucleoid structure. Further analysis reveals that Shield can be divided into four subtypes, three of which contain additional components that in some cases can modulate the activity of ShdA and/or provide additional lines of phage defence. Collectively, our results identify a new player within the *Pseudomonas* bacterial immunity arsenal that displays a novel mechanism of protection, and reveals a surprising role of RmuC domains in phage defence.

**SIGNIFICANCE:** The evolutionary pressure exerted by bacteriophages has driven bacteria to acquire numerous defense systems. Recent studies have highlighted the extraordinary diversity of these systems, uncovering exciting links between bacterial and eukaryotic immunity. Here we describe a novel anti-phage system, named Shield, found within *Pseudomonas* species. We identify several Shield subtypes, all harboring the same core component, and describe its mode of action. The growing instance of multidrug-resistant bacterial infections urgently requires the development of alternative treatments. Phage therapy is a particularly pertinent approach to treat multi-drug resistant *Pseudomonas aeruginosa* strains causing severe lung infection in cystic fibrosis patients. A detailed understanding of bacterial immunity and phage counter-strategies is an essential step to underpin the rational design of phage therapy to fight disease.

## INTRODUCTION

In response to continuous predation from bacteriophages (phages), bacteria have evolved numerous defense systems that, collectively, can be considered to provide ‘bacterial immunity’ (1–4). Historically, the best described phage defense systems are the restriction-modification (RM) systems, which represent an example of bacterial innate immunity. Here the modification unit covalently modifies host DNA whilst the restriction component recognises specific sequence patterns on unmodified invading DNA to mediate its degradation (5). The more recently discovered CRISPR-Cas systems represent the first example of bacterial adaptive immunity wherein guide RNAs, specific for invading phage DNA sequences, direct effector nucleases to foreign nucleic acids (6). Understanding the mechanism of action of these systems has underpinned their utility as tools that revolutionised the gene editing and molecular biology fields. RM and CRISPR-Cas systems are usually considered part of the first line of defense, as their defense strategy is to swiftly remove invading phage DNA (7). Conversely, several more recently-discovered anti-phage systems represent a second line of defense, where their mode of action induces programmed cell death of the infected cells (7). This strategy is broadly defined as abortive infection (Abi) and represents a possible altruistic behaviour wherein infected cells induce their own premature death in order to prevent release of mature phage particles within the population (7). The division between ‘first’ and ‘second’ lines of defense was recently shown to be less defined, with some instances of CRISPR-Cas subtypes also inducing bacterial stasis or death (8, 9).

A defining trait of known anti-phage systems is that they are frequently encoded together at genomic hotspots, known as ‘defense islands’ (8). This concept has been exploited in recent years to identify novel defense loci encoded close to known systems, revealing numerous additional anti-phage systems that were never described before (10–15). This has uncovered further innovative defensive strategies including NAD+ depletion, mediated by the Thoeris and Defense-associated sirtuin (DSR) systems (16, 17) and depletion of the cellular dNTP pool (18, 19). Depletion of cellular dNTPs is a conserved antiviral strategy also found in eukaryotes, demonstrating a clear link between prokaryotic and eukaryotic immunity (20). Indeed, a striking characteristic that has emerged from recent studies is the evolutionary relatedness between many anti-phage systems and innate immune mechanisms in plants and animals (12, 13, 18, 21–25). Additional examples of bacterial anti-phage defense systems that are ancestors of eukaryotic immunity are represented by the viperins, gasdermins, NLR-related anti-phage systems, Toll/IL-1 receptor (TIR) domains (carried by the Thoeris system) and the bacterial cyclic oligonucleotide-based signalling system (CBASS) (12, 13, 18, 21–25). However, despite these recent discoveries, the defense mechanisms adopted by the majority of the newly-identified anti-phage systems remain unknown. Furthermore, these recent approaches have highlighted that many more genes involved in bacterial immunity still await discovery.

In this study we report the identification of a previously uncharacterised phage defense system that we have termed Shield. The system, which is encoded across *Pseudomonas* species, has an RmuC domain-containing protein, ShdA, as its active component. ShdA exhibits DNA-degrading activity *in vitro,* whereas expression *in vivo* leads to a remodelling of the host nucleoid structure as part of the defense mechanism. While ShdA alone is sufficient to confer phage defense, Shield occurs as four distinct subtypes. In three of these, additional components are found. We demonstrate that additional components of two subtypes regulate ShdA activity. Collectively, our study uncovers the previously uncharacterised role of RmuC domains in bacterial immunity and describes the RmuC-mediated mechanism of phage inhibition.

## RESULTS

### Genetic identification of a candidate novel anti-phage system

Previous studies have demonstrated that bacterial defense systems cluster together in chromosomal hotspots, defined as ‘defense islands’ and that their mobilisation is dependent on mobile genetic elements (4, 25, 26). Genetic neighborhood analysis of known anti-phage systems has allowed the systematic discovery of many novel defense systems (10, 11, 13, 14). Following the same principle, we aimed to identify bacterial operons, situated in the context of defense islands, whose role has not yet been associated with defense against invasion of foreign DNA.

*Pseudomonas aeruginosa* is among the species that have been reported to encode many distinct anti-phage systems (27). To identify new candidate anti-phage systems in this organism and across the genus we downloaded all available genome sequences of *Pseudomonas* species from the Refseq database to the ‘Scaffold’ assembly level (ftp://ftp.ncbi.nlm.nih.gov/genomes/ASSEMBLY_REPORTS/assembly_summary_refseq.txt, as of 11th June 2022) and used the defense-finder tool to identify known anti-phage systems (27) (Table S1). Next, we manually screened for flanking genes or operons that associated with the defense systems identified in Table S1 and whose function was either unknown (genes annotated to encode hypothetical proteins), associated with interaction/degradation of nucleic acids, or that could mediate cell lysis. Our search consistently revealed a gene that was encoded next to several known anti-phage systems (Figure 1a). Using the protein encoded by *P. aeruginosa* NCTC 11442/ATCC 33350 (assembly ID: GCF_001420205.1, locus tag: AN400_RS26690) we performed functional predictions with hmmscan against a PFAM database (26), identifying that the C-terminal region of AN400_RS26690 contains a PF02646 domain, typical of RmuC proteins (Figure 1b). RmuC is involved in recombination and is associated with regulating the rate of inversion events at short-inverted repeats (28). Additionally, AN400_RS26690 harbors a predicted trans-membrane domain (TMH) at its N-terminus (Figure 1b). Based on results presented later, we subsequently renamed this protein ShdA.

**Figure 1:**
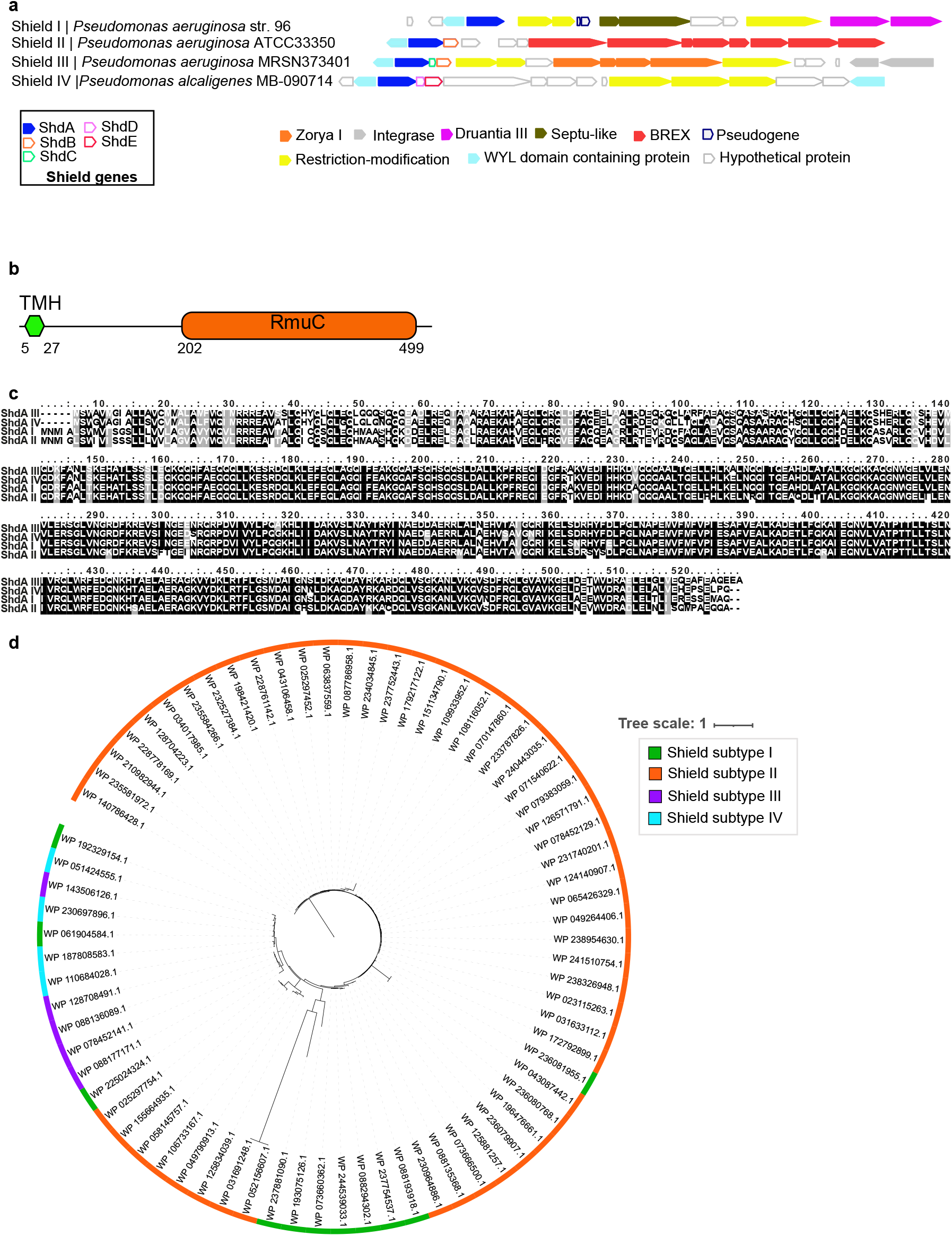
Identification of a novel anti-phage system. **(a)** Schematic of the genomic neighbourhood of Shield systems in representative genomes. Known defense genes were predicted using PFAM and defense-finder (1). ShdA is in blue and other Shield partners are represented with coloured outline, as indicated. The full set of Shield subtypes is shown in Figure S2 and known anti-phage systems annotations are reported in Table S6. **(b)** Schematic representation of the predicted domain organisation of the candidate defence protein, ShdA. **(c)** Representative alignments of ShdA homologues across Shield subtypes. One representative ShdA homologue was chosen for each Shield subtype and aligned using MUSCLE. A full alignment, involving all homologues, is shown in Figure S2. **(d)** Phylogenetic tree based on ShdA homologues. Coloured blocks indicate ShdA homologues belonging to each Shield subtype.

Subsequently, to extend the search and detect a wider diversity of homologues, we built a Hidden Markov Model (HMM) model, including RmuC-like proteins from several species. We then queried a local Refseq of all bacterial genomes to the ‘contig’ assembly level (ftp://ftp.ncbi.nlm.nih.gov/genomes/ASSEMBLY_REPORTS/assembly_summary_refseq.txt, as of 11th June 2022). The dataset was manually curated to include only those homologues found within defense islands (thus excluding housekeeping RmuC) (Figure 1a, Figure S1). This resulted in ~70 unique protein identifiers, and flanking gene (FlaGs) analysis (29) was subsequently used to define the genomic neighborhood of the *shdA* genes (Figure 1a, Figure S1, Table S2, Table S3). From this we noted that although in some instances *shdA* could be found as an orphan gene, in other instances it co-occurred with subsets of additional genes which were always encoded downstream (Figure 1a, Figure S1, Table S2). This allowed us to classify the predicted anti-phage system, which we have named ‘Shield’ into four subtypes (Figure 1a, Table 1).

**Table 1.** Summary of predicted protein components for each Shield subtype.

| Shield subtype | System components |
| --- | --- |
| Shield I | ShdA I |
| Shield II | ShdA II, ShdB II |
| Shield III | ShdA III, ShdB III, ShdC |
| Shield IV | ShdA IV, ShdD, ShdE |

Where ShdA was encoded alone, we designated this Shield I. Of the other three candidate systems, Shield subtype II is the most widely distributed (Figure 1, Figure S1 and Table S6) and we renamed its partner gene *shdB.* We named the other partner genes *shdC* - *shdE* according to the order they appeared in the FlaGs schematic representation of Shield subsystems (Figure 1a, Table 1). A BLAST search with strict query coverage and sequence similarity parameters (80-100% and 70-100%, respectively) revealed that ShdC-ShdE proteins are predominantly found in association with Shield subsystems.

The 15 proteins encoded directly upstream and downstream of *shdA* were annotated using defense-finder and PFAM predictions (Figure 1a, Figure S1, Table S3-S5), confirming that Shield subtypes localize adjacent to known anti-phage systems (Figure 1a, Figure S1, Table S3 - S5). They also often associate with WYL-domain containing proteins, transcriptional regulators that are enriched in phage defense islands (3, 30, 31). Furthermore, the genome neighborhood also encodes several viral proteins and integrases, in agreement with recent reports that mobile genetic elements (MGE) represent primary carriers of defense islands (14, 26, 32). These findings further support the involvement of the Shield systems in bacterial immunity.

Alignment of ShdA homologues reveals a high degree of sequence identity, particularly towards the C-terminus (Figure 1c, Figure S2). A phylogenetic tree (based on the alignment in Figure S2), reveals that although Shield subtype III is more similar to Shield subtype II in terms of gene composition (both having a ShdA and ShdB homologue), ShdA III homologues cluster closer to homologues from subtype I and IV (Figure 1d), whilst some ShdA I proteins are found in-between ShdA II branches (Figure 1d). Finally, despite the relaxed parameters used in search of RmuC homologues, Shield subtypes were, to date, only found in *Pseudomonas spp* (Table S6).

Within our ShdA homologues hits, we also found ~70 homologues embedded within DISARM-like operons (Figure S3a, S4, Table S7). DISARM-associated ShdA proteins exhibit a longer sequence length, with an additional sequence stretch at the N-terminus (Figure S5). Phylogenetic analysis shows that DISARM-related ShdA homologues cluster separately from those of Shield, suggesting ShdA homologues associated to Shield and DISARM have diverged early (Figure S3b). Interestingly, the taxonomic distribution of DISARM-associated ShdA proteins is also exclusively limited to *Pseudomonas spp.* (Table S8).

### ShdA is the core anti-phage defense module in Shield subtypes

Since ShdA is the sole component of Shield I, we reasoned that this protein might represent the core ‘defense’ module of Shield systems. To investigate this, we cloned representative examples of *shdAI* - *shdAIV* under control of the arabinose-inducible promoter in plasmid pBAD18. We subsequently introduced these constructs into *Escherichia coli* MG1655 and tested their ability to protect *E. coli* growing in liquid culture from lysis by phages ϕSipho and ϕAlma (Figure 2).

**Figure 2:**
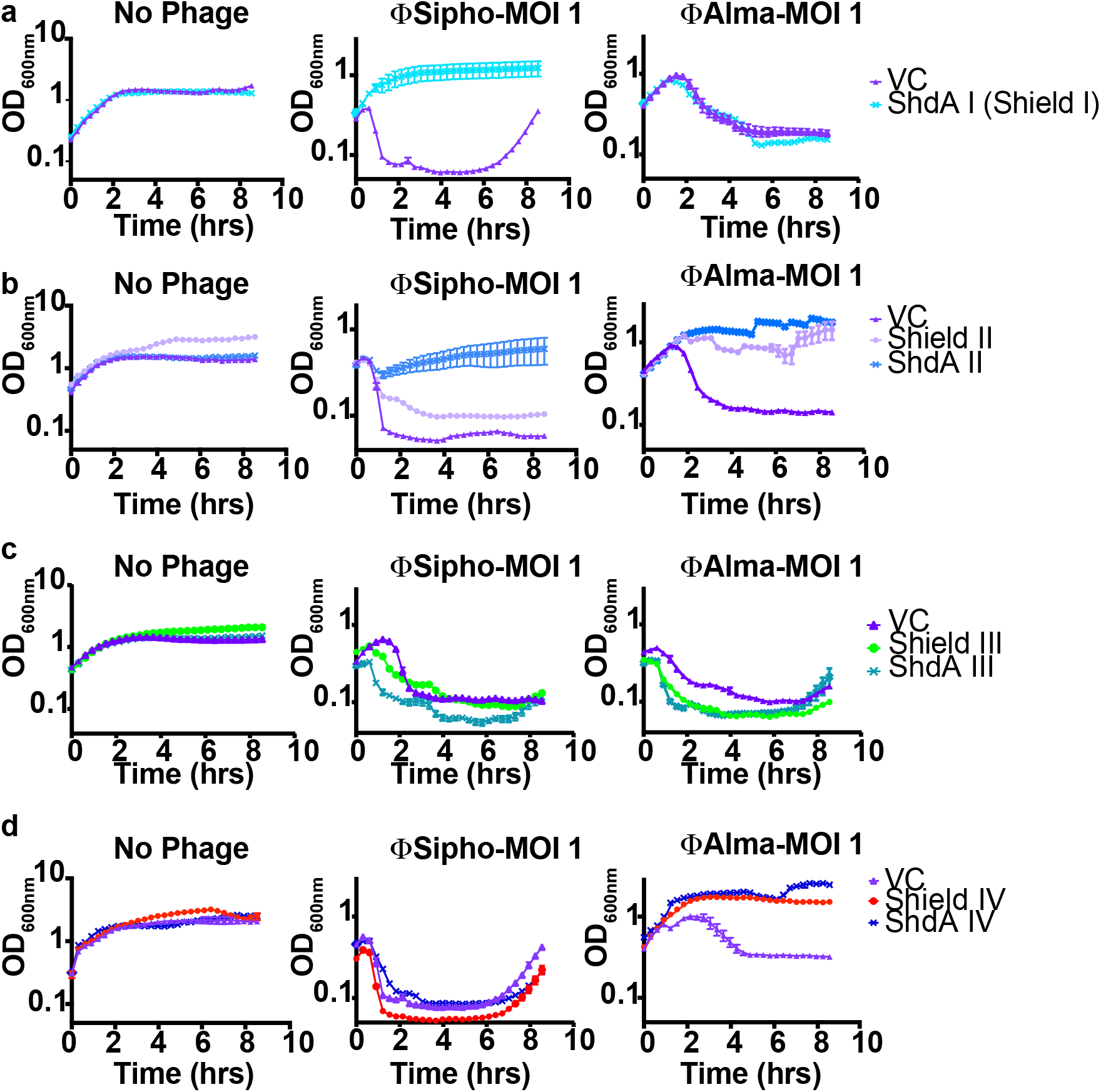
Shield I, Shield II and Shield IV prevent phage infection. Growth curves of *E. coli* MG1655 carrying empty pBAD18 (VC) or the same plasmid encoding **(a)** ShdA I (the sole component of the Shield I system), **(b)** the Shield II system or ShdA II only, **(c)** Shield IV or ShdA IV only, and **(d)** Shield III or ShdA III only. Strains were grown in LB medium supplemented with 0.2 % L-arabinose and phages ϕSipho or ϕAlma were added at the start of the growth curve at a MOI = 1. Points show mean +/- SEM (n = 3 biological replicates).

We observed that Shield I, which only comprises the ShdA I module, confers robust protection against the phage ϕSipho (Figure 3a), but was ineffective against ϕAlma. Similarly, ShdA II and ShdA IV also provided defense against phage-mediated lysis (Figure 2b, 2d), with ShdA II protecting against the action of both ϕSipho and ϕAlma, and ShdA IV against ϕAlma only. We also tested the ability of the full Shield II and Shield IV systems to provide protection, and noted that in each case the cognate ShdA elicited at least similar levels of protection as the full systems (Figure 2b, 2d). Conversely, ShdA III and Shield III were ineffective against both phages (Figure 2c). We conclude that at least three of the Shield subtypes are involved in bacterial immunity, and that the defense phenotype is dependent on the conserved component ShdA. We also conclude that the different subtypes we tested show phage-specific patterns of protection.

**Figure 3:**
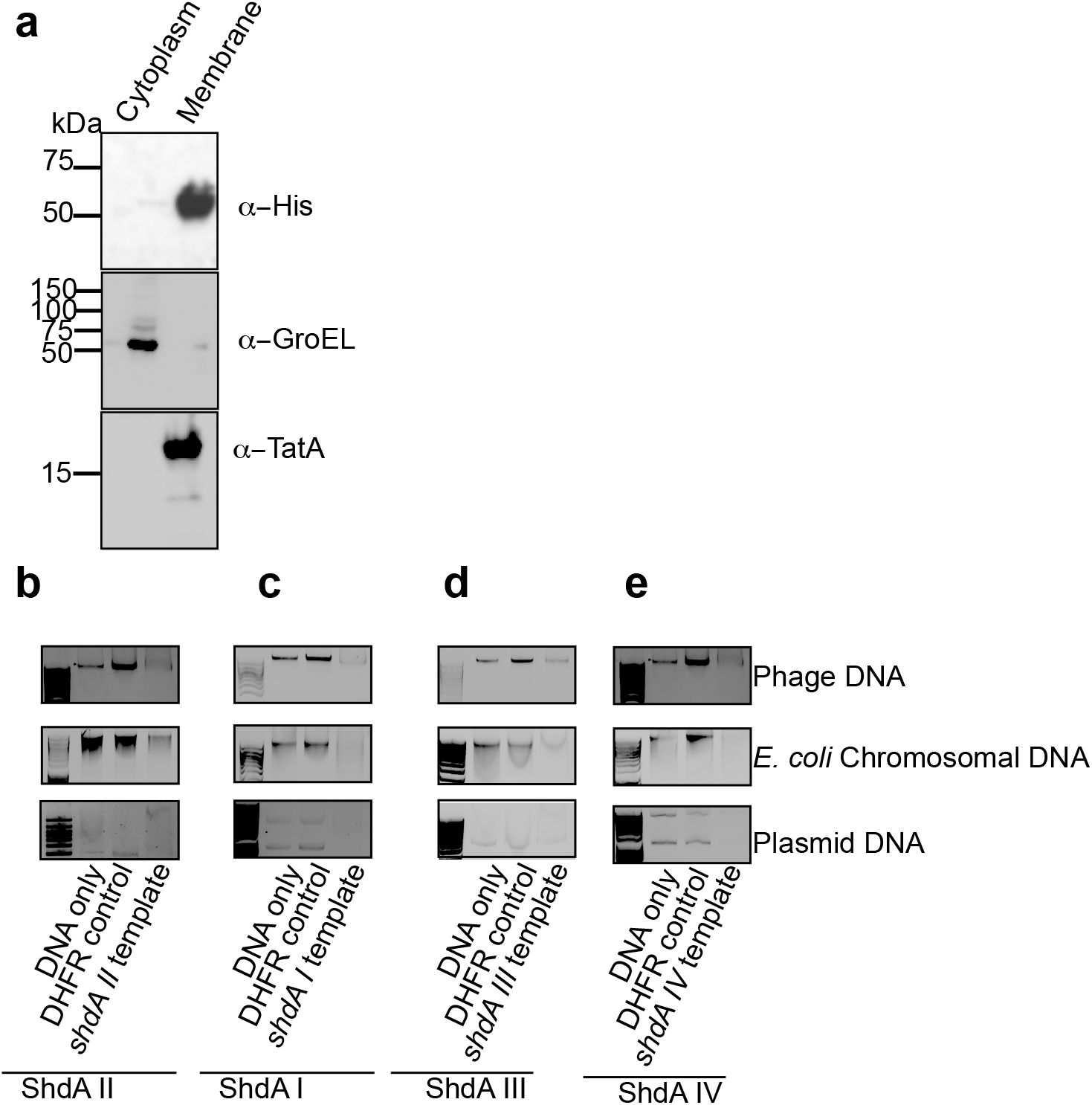
ShdA homologues exhibit nuclease activity *in vitro.* **(a)** Cells carrying plasmid pBAD18 encoding ShdA II-His6 were grown for 5 hrs in the presence of 0.2 % L-arabinose. Cells were fractionated to produce soluble and membrane samples and analysed by immunoblot with antibodies to the His6 tag, GroEL (cytoplasmic control) and TatA (membrane control). **(b-e)** *In vitro* DNAse activity assays using **(b)** ShdA II_138-524_ **(c)** ShdA I_140-526_ **(d)** ShdA III_135-523_ and **(e)** ShdA IV_135-521_. ShdA proteins and DHFR were synthesised using the cell-free PURExpress kit (NEB). DNAse activity was tested against 10 ng of input DNA. DNA types tested were phage DNA, *E. coli* MG1655 chromosomal DNA and plasmid (pSG483) DNA. For ShdA I_140-526_, ShdA III_135-523_ and ShdA II_138-524_ phage DNA was from ϕSipho. For ShdA IV_135-521_ phage DNA was from ϕAlma.

### ShdA is a membrane protein that has nuclease activity *in vitro*

Most of ShdA homologues we identified are predicted to have a TMH domain at their N-terminus (Table S9). To confirm that this prediction is correct, we fractionated *E. coli* cells producing C-terminally His-tagged ShdA II. Western blot analysis revealed that the protein was only detected in the membrane fraction (Figure 3a). We then asked the question whether membrane-localization of ShdA is essential for its anti-phage activity. To this end we attempted to clone truncated *shdA* genes lacking the 5’ regions encoding the membrane anchor. However, all our efforts were unsuccessful, even when we used tightly repressible vectors, strongly suggesting that the encoded proteins were toxic to *E. coli*. Thus, it seems likely that anchoring of ShdA to the membrane modulates its toxicity.

It was previously shown that *E. coli* RmuC displays a restriction endonuclease-like fold and is predicted to regulate genome inversion events through DNA cleavage (33). We therefore next sought to test whether the RmuC domain in ShdA has nuclease activity. As we were unable to clone the truncated *shdA* alleles, we used cell-free synthesis to generate the proteins. Following cell-free synthesis using a template for ShdA II_138-524_, the products caused degradation of phage ϕSipho DNA, *E. coli* chromosomal DNA and plasmid DNA (Figure 3b). No degradation was observed when using products generated with the kit-provided template for the control protein dihydrofolate reductase (DHFR) (Figure 3b).

To confirm whether this was a common feature of ShdA proteins, we also tested the ability of *in vitro* synthesised ShdA I_140-526_, ShdA III_135-523_ and ShdA IV_135-521_ to degrade DNA. Whilst the yield of each was low, the products all showed DNase activity (Figure 3c-e), confirming that DNA degradation is a common feature of ShdA proteins. Interestingly, although we were not able to identify any phages in our collection that were sensitive to Shield III, the ShdA III protein was capable of partially degrading DNA from phage ϕSipho. This suggests that ShdA III is also likely to be active in phage defense, but that our phage collection does not contain representative phage that are sensitive to its activity.

### Shield II and ShdA II mode of action involves reduction of phage burden

To investigate the mode of action of Shield in phage defense *in vivo,* we focused on Shield II from *P. aeruginosa* NCTC 11442/ATCC 33350 (assembly ID: GCF_001420205.1), as this represents the most common Shield subtype (Table S6).

We first tested the ability of ShdA II and Shield II to provide protection against a suite of phages. We found, through calculation of fold protection, that ShdA II conferred resistance to several different phages (Figure 4a). The same phages are also susceptible to Shield II-mediated defence as indicated by a decrease of efficiency of plating (EOP) in Figure 4b. Subsequently, we tested the EOP of strains producing either the full Shield II system or ShdA II or ShdB II alone when infected with phages ϕSipho, ϕTB34, ϕAlma, ϕNR1 or ϕBaz. We observed that both Shield II and ShdA II conferred a decrease in EOP in all cases (Figure 4c), and that they decreased the burst size of ϕSipho (Figure 4d). Additionally, as *in vitro* analysis of ShdA revealed the conserved ability to degrade plasmid DNA (Figure 3), we tested whether Shield II/ShdA II could affect the efficiency of plasmid DNA acquisition during transformation. Indeed, both Shield II and ShdA II decreased transformation efficiency (Figure 4e).

**Figure 4:**
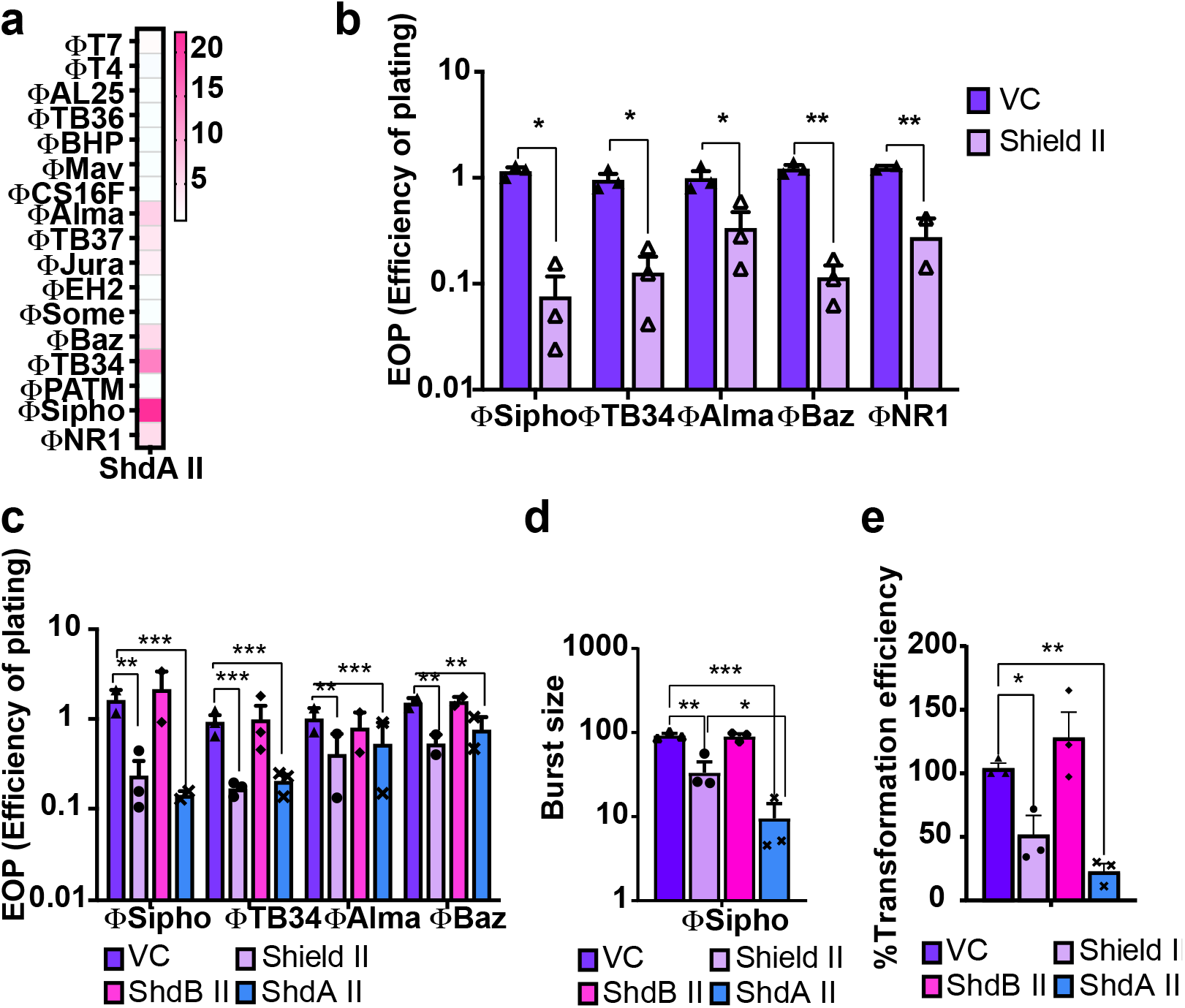
Characterisation of the anti-phage activity of Shield subtype II. **(a)** Evaluation of ShdA II fold protection against a suite of phages. Fold protection was calculated by comparing the number of phage plaques formed on *E. coli* MG1655 producing ShdA II with the number of plaques generated from the same strain harboring empty vector. **(b)** Efficiency of plating (EOP) measurement for *E. coli* MG1655 carrying empty vector (VC, pBAD18) or the same plasmid encoding Shield subtype II when infected with the indicated phages. Points show mean +/- SEM (n = 3 biological replicates). **(c)** Efficiency of plating (EOP) measurement for *E. coli* MG1655 carrying empty vector (VC, pBAD18) or the same plasmid encoding the Shield II system, ShdA II only or ShdB II only when challenged with phages ϕSipho, ϕTB34, ϕAlma, ϕBaz and ϕNR1. Points show mean +/- SEM (n = 3 biological replicates) except for phages ϕSipho and ϕTB34 where n = 4 biological replicates **(d)** Average burst size assessment for the same strain and plasmid combinations as (c) following infection with ϕSipho at MOI 0.1. Points show mean +/- SEM (n = 3 biological replicates) **(e)** Evaluation of transformation efficiency of the same strain and plasmid combinations as shown in panel (c) with plasmid DNA. For all panels, 0.2% L-arabinose was added at time zero to induce expression of the encoded genes in pBAD18. Points show mean +/- SEM (n = 3 biological replicates). Statistical analysis for panel **b** was performed with GraphPad applying unpaired student t test. No significance was detected, unless indicated (*p≤0.05). For panels **c-e**statistical relevance was measured using one-way ANOVA with Dunnett’s multiple comparison test. No significance was detected, unless indicated (*p≤0.05).

Next, we investigated Shield II and ShdA II-mediated defense in liquid culture against ϕSipho or ϕTB34. Cells harboring Shield II or ShdA II showed better survival than cells carrying empty vector (VC) or ShdB II only, especially at lower MOI values. Additionally, ShdA II-expressing cells showed a better growth rate than Shield II-harboring cells, during phage infection (Figure 5a).

**Figure 5:**
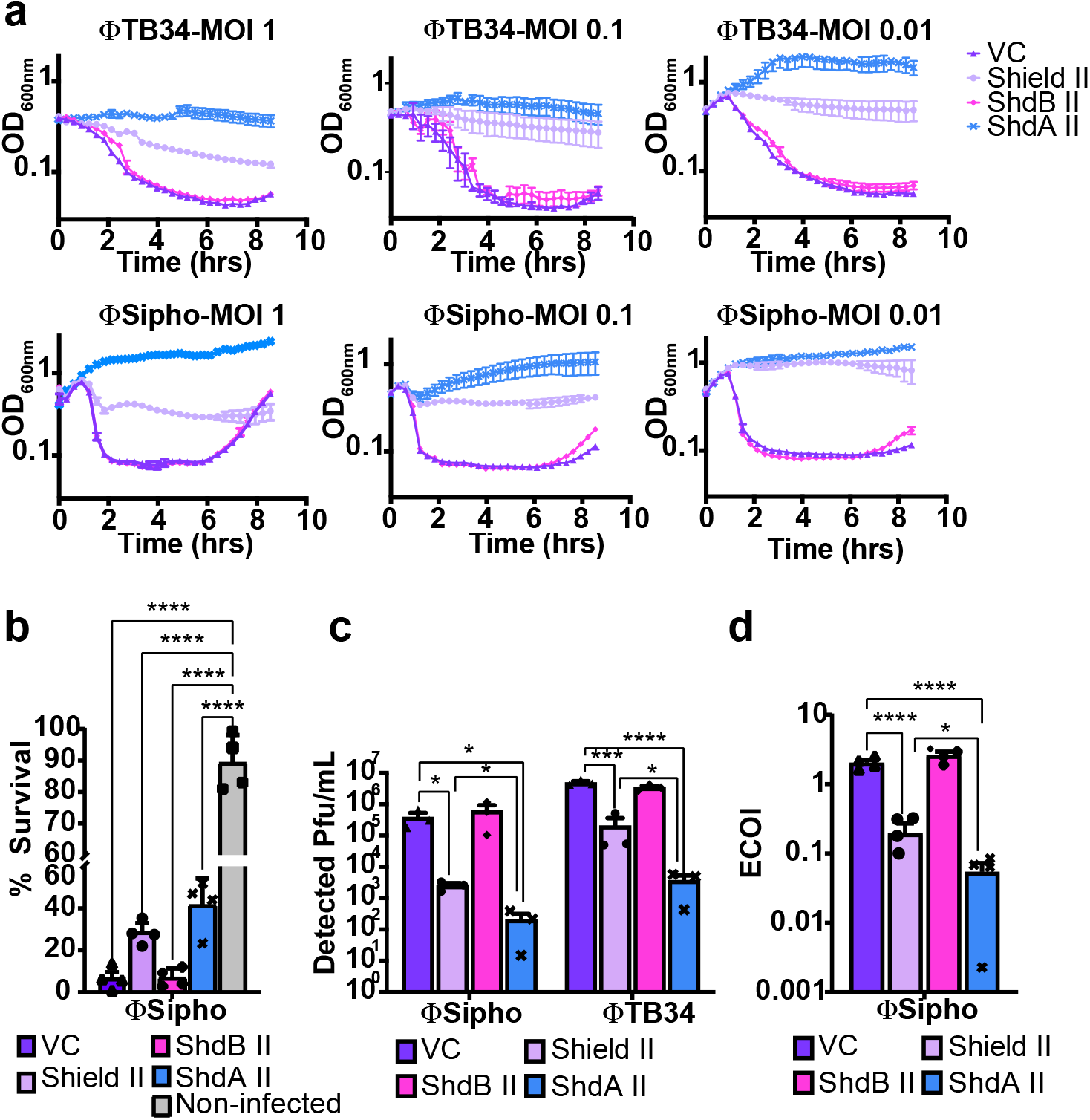
ShdA II reduces the phage burden through a mechanism reminiscent of abortive infection. **(a)** Growth curves of *E. coli* MG1655 carrying pBAD 18 (VC,) or the same plasmid encoding Shield II, ShdA II only or ShdB II only. Strains were grown in LB supplemented with 0.2 % L-arabinose and infected at time zero with MOI = 1, MOI = 0.1, MOI = 0.01 of ϕSipho and ϕTB34, as detailed in Methods. Evaluation of **(b)** Cell survival, **(c)** phage replication and **(d)** efficiency of centre of infection (ECOI) for *E. coli* MG1655 carrying pBAD 18 (VC) or the same plasmid encoding Shield II, ShdA II only or ShdB II only when challenged with phage ϕSipho at MOI = 0.1. For each assay, strains carrying the indicated plasmid were grown in LB supplemented with 0.2% L-arabinose and assays were performed as described in Material and Methods. For all panels, points show mean +/- SEM (n = 3 biological replicates). Statistical relevance was measured using one-way ANOVA with Dunnett’s multiple comparison test. No significance was detected unless indicated (*p≤0.05).

To better determine how ShdA II mediates phage defense, we measured cell survival following infection with ϕSipho (Figure 5b). Cells harboring Shield II or ShdA II both displayed a slightly increased survival rate, compared to cells containing vector only, or producing ShdB II alone (Figure 5b). In both cases, survival was not complete when compared to non-infected controls, which is normally a hallmark of abortive infection (7). Consistently, quantification of phage produced during infection, as previously reported (34), showed that in the presence of Shield II or ShdA II both ϕSipho and ϕTB34 replication was hindered (Figure 5c). Nevertheless, the inhibition was more potent in ShdA II-harboring cells compared to cells expressing the entire Shield II system (Figure 5c). Additionally, when we measured the Efficiency of Centre Of Infection (ECOI), we observed that in presence of ShdA II a fewer number of infective centres were released compared to the full Shield II (Figure 5d). While liquid culture assays, cell survival assays and phage replication measurements (Figure 4 and 5) suggest that Shield II mode of action could include a mechanism akin to abortive infection, the phenotypes observed could be partially due to incomplete clearance of the tested phage’s infection provided by Shield II (Figure 5c-d).

From the experiments evaluating the cell survival in liquid cultures, phage replication, burst size, and ECOI measurements of strains carrying Shield II and ShdA II, we consistently noted that presence of ShdA II alone reduces the amount of released phage particles to a greater extent than that observed for cells harboring Shield II. Taken together these results suggest that ShdB II may negatively regulate the activity of ShdA II.

### ShdA induces bacterial nucleoid re-arrangement that is modulated by ShdB

The *in vitro* nuclease assays (Figure 3b-e) indicated that the RmuC domain of ShdA has DNase activity that is active against chromosomal DNA. To determine whether cells producing ShdA II were anucleate or exhibited other types of aberrant cellular organisation such as changes in degree of condensation, we examined them using the DNA-specific stain 4’,6-diamidino-2-phenylindole (DAPI) and fluorescence microscopy. We observed that over a timecourse of ShdA II induction, *E. coli* chromosomal DNA was not detectably depleted/degraded, indicated by the absence of anucleate cells. However, the bacterial nucleoid underwent drastic spatial rearrangement within the cell (Figure 6a). After 2 hr of induction, the nucleoid of ShdA II-expressing cells adopted a distinct morphology and cellular localisation, with an increasingly patchy and peripheral distribution pattern (Figure 6a-b). This data suggests that ShdA II likely able to interact with chromosomal DNA and recruit it to the cell periphery. Cells producing Shield II showed a less extensive redistribution of DAPI fluorescence, again consistent with a role for ShdB II in modulating the function of ShdA II.

**Figure 6:**
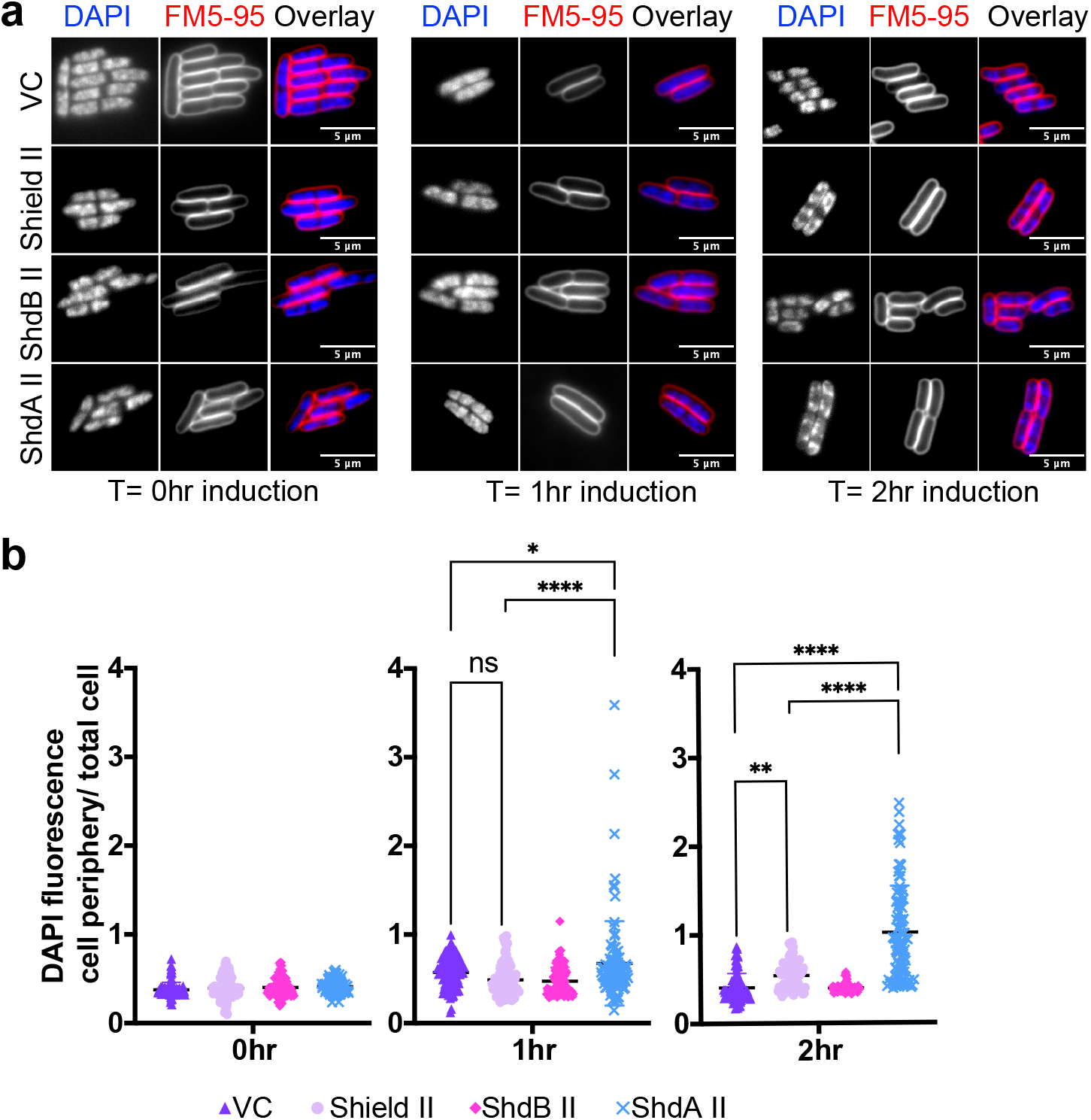
ShdA alters bacterial nucleoid morphology. **(a)** *E. coli* MG1655 harboring pBAD18 (pBAD18) or the same plasmid encoding Shield II, ShdA II only or ShdB II only were cultured in LB medium containing 0.2% L-arabinose for 2 hours. At t=0 hours, t=1 hour and t=2 hours an aliquot of each culture was removed and stained with DAPI (for DNA visualisation) and FM 5-95 dye (to stain cell periphery/ outer membranes). Cells were subsequently imaged using fluorescence microscopy **(b)** The ratio between DAPI fluorescence intensity at the cell periphery and the whole cell was quantified to assess ShdA II-mediated recruitment of DNA at the cell periphery for strains in **(a)**. Points show mean +/- SEM (n = 100 cells)

### ShdB is a probable peptidase that regulates the cellular level of ShdA

To investigate the role of ShdB in the Shield II system we generated a high confidence structural model using AlphaFold (Figure S6a-c) and used it to search the Dali server (35, 36). This predicted structural similarity to the metallopeptidase M15 family and to several carboxypeptidases (Figure S6a-c and Table S10). While M15 family proteins are normally involved in peptidoglycan biosynthesis and turnover, ShdB harbors no detectable signal peptide that would direct it to the periplasm. Furthermore, when we provided ShdB II with the signal peptide from OmpA, a substrate of the *E. coli* Sec pathway, we observed no effect on the growth of *E. coli,* suggesting that the protein is unlikely to cleave peptidoglycan (Figure S7).

Interestingly, some toxin-antitoxin systems have antitoxins with predicted metallopeptidase activity that exert their effect through degradation of the toxin partner (13, 37, 38). We therefore reasoned that ShdB may similarly be a protease whose specific substrate is ShdA. To explore this further we first tested the ability for ShdA II and ShdB II to interact using the bacterial two hybrid assay (BTH) (39). As shown in Figure 7a, we observed an interaction between ShdA II and ShdB II when the T25 fragment of the adenylate cyclase was fused to ShdA II and the UT18 fragment to ShdB II (Figure 7a). Furthermore, when we modified *shdAII* in the constructs that were used to assess anti-phage activity to produce a protein with a C-terminal hexahistidine epitope, a drastic decrease in ShdA-His6 protein levels was observed when coproduced with ShdB II (Figure 7b). Taken together our results are consistent with the hypothesis that ShdB II regulates the activity of ShdA II through targeted proteolysis.

**Figure 7:**
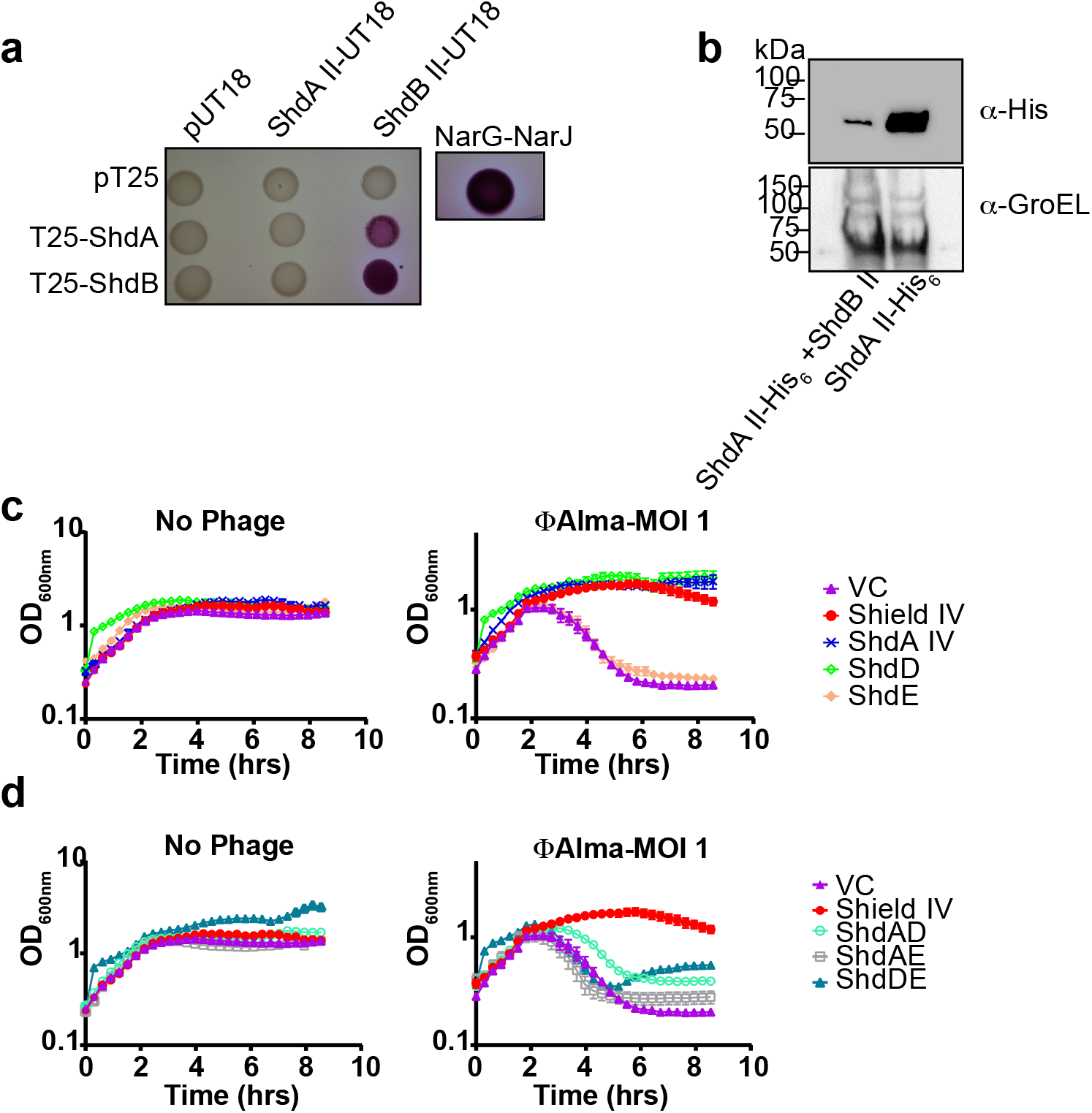
ShdA-associated components in Shield subtypes regulate ShdA activity and/or provide additional defence modules. **(a)** Analysis of *E. coli* BTH101 carrying combinations of ShdA II and ShdB II, when cloned in either pUT18 or pT25 vectors as indicated. The empty pUT18 or pT25 vectors were used as negative controls, while the interaction between NarG and NarJ proteins was employed as a positive control (2). **(b)** ShdA II-His6 or ShdA II-His6 + ShdB II were expressed from an arabinose-inducible plasmid pBAD18. and ShdA II-His6 levels were assessed by western blot analysis. GroEL was used as loading control. **(c)** Growth in liquid LB supplemented with 0.2 % L-arabinose of *E. coli* MG1655 carrying (VC, pBAD18) or plasmids directing the expression of the Shield IV system (Shield IV), ShdAIV only (ShdAIV), ShdD only (ShdD), ShdE only (ShdE). Strains were infected with MOI = 1 of ϕAlma as described in Material and Methods. Point show mean +/- SEM (n= 3 biological replicates). **(d)** *E. coli* MG1655 carrying (VC, pBAD18) or plasmids directing the expression of the Shield IV system (Shield IV), the ShdA IV and ShdD pair (ShdAD), ShdA IV and ShdE pair (ShdAE), or the ShdA and ShdE pair (ShdDE) were grown in LB medium supplemented with 0.2 % L-arabinose and infected with MOI = 1 of ϕAlma as indicated in Methods. Point show mean +/- SEM (n= 3 biological replicates).

### Accessory Shield components also regulate ShdA mediated defense

Finally, we sought to determine whether a similar regulatory mechanism is found for Shield IV, a subtype that contains the ShdD and ShdE components. Neither of these proteins have any identifiable domains or predicted functions, and share no homology with ShdB. Using phage ϕAlma, we found that ShdD alone was, like ShdA IV, capable of providing protection (Figure 7c), indicating that two phage defense modules are part of Shd IV. However, while the full Shield IV system was competent to mediate resistance to ϕAlma infection, none of the pairwise combinations conferred any significant protection, suggesting that unlike Shield II, the regulation and fine-tuning of Shield IV anti-phage activity is more complex (Figure 7d).

## DISCUSSION

Here we report the discovery of a previously uncharacterised anti-phage system, Shield, demonstrating that it reduces the release of mature viral particles during phage infection by abortive infection (Figure 4–5). The basic defense module of Shield is ShdA, a membranebound protein with a cytoplasmically-located RmuC domain (Figure 2–3). When produced *in vitro,* ShdA RmuC domains have non-specific (endo)nuclease activity and can degrade phage, plasmid and chromosomal DNA (Figure 2). Our inability to clone or express truncated forms of ShdA lacking the membrane anchor strongly suggests that cytoplasmic ShdA is genotoxic and implies that membrane attachment is key to Shield function (Figure 2). In agreement with this, full length ShdA is noticeably less toxic when produced *in vivo.* However, full length ShdA does trigger striking recruitment of chromosomal DNA to the cell periphery *in vivo* (Figure 6). This implies that some feature of the membrane, for example steric hindrance, regulates ShdA DNase activity. It is worth highlighting that while the observed recruitment of DNA to the membrane is indicative of strong ShdA-DNA interactions, the extent of this recruitment is likely exacerbated by high expression levels. Certain other phage defense systems also rely on nuclease activity, for example NucC from the CBASS system, to bring about abortive infection through degradation of chromosomal DNA (40). However, examples of anti-phage defense mediated by a co-opted RmuC domain were not reported before.

While ShdA is clearly a central component of Shield-mediated defense, Shield is found as at least four distinct subtypes, with three of them having additional component/s. Two of the three, Shield II and Shield III, contain the ShdB protein. ShdB is a negative regulator of ShdA,and is structurally related to metallopeptidases. Our experimental findings indicate that ShdB interacts with ShdA and likely degrades it to modulate the cellular level of ShdA. Another intriguing, though speculative, possibility is that ShdB can release ShdA from the membrane upon phage infection, thereby activating ShdA’s nuclease activity. The Shield IV system comprises ShdA and two further components, ShdD and ShdE. Like ShdB, both ShdD and ShdE negatively regulate ShdA activity, however the situation is more complex because ShdD can also act as a phage defense module when expressed in isolation. Further work would be required to elucidate the roles of ShdD and ShdE in Shield IV-mediated defense.

Figure 8 outlines models that may account for the activity of the Shield I and Shield II systems. In each case, under non-infection conditions, ShdA is present in the membrane and interacts with, but does not degrade, chromosomal DNA. We suggest that the initial stages of phage infection result in changes at the cell envelope, for example phage invasion triggering membrane depolarisation or bulging of the cytoplasmic membrane associated with tail tube fusion (41–43). Associated changes in membrane physical properties may activate the nuclease activity of membrane-bound ShdA through conformational change or proteolytic cleavage, resulting in chromosomal and phage DNA degradation and cell death before phage replication can occur. For the Shield II system, we propose that ShdB maintains ShdA at a less active state. We suggest that while this level may potentially be sufficient to provide some degree of protection, the ShdB component provides a second checkpoint - for example titration of ShdB away from ShdA by a phage component would allow ShdA to accumulate to higher levels ensuring protection. Alternatively, partial proteolytic cleavage of ShdA by ShdB could trigger activation of ShdA-dependent DNA degradation through release from the membrane.

**Figure 8:**
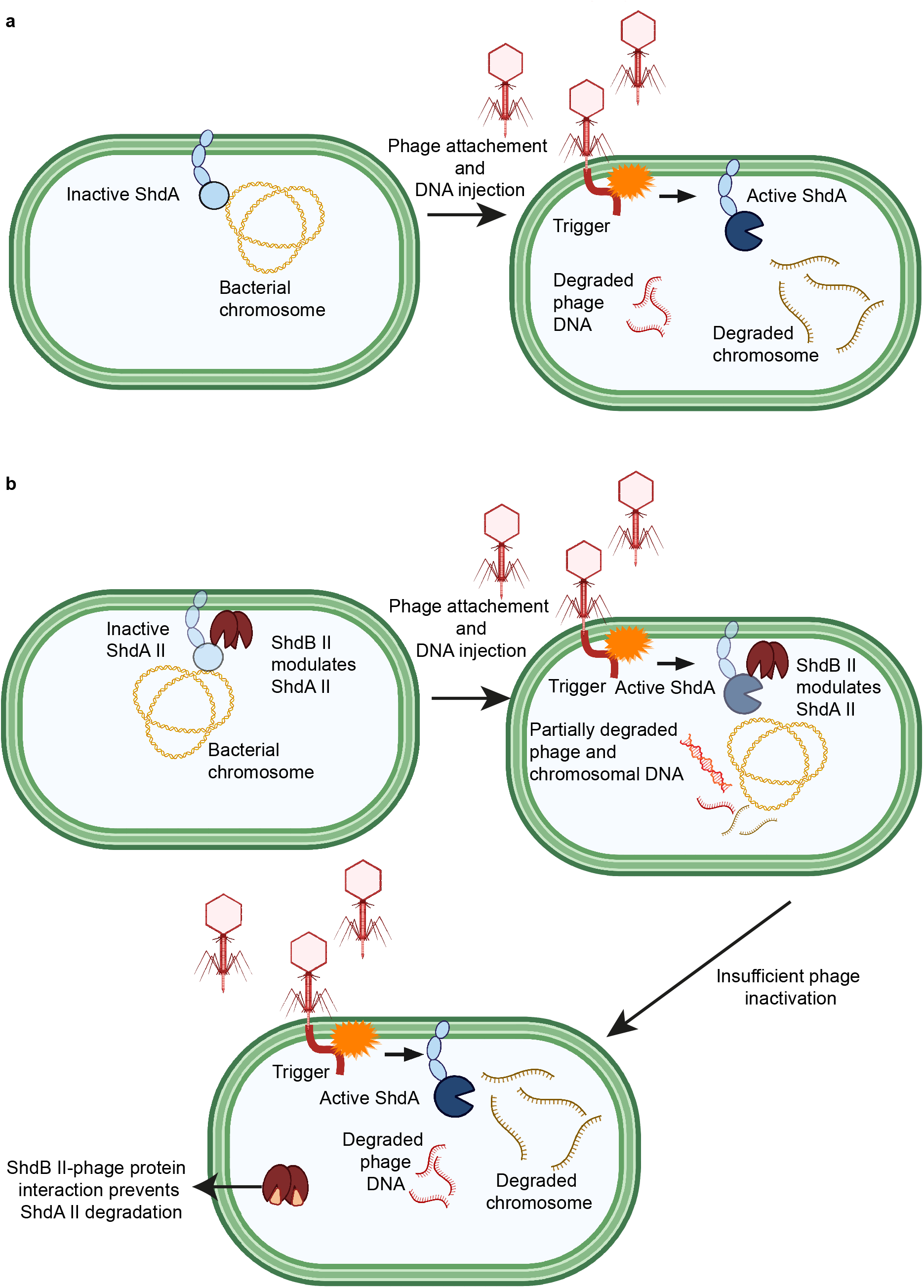
Model for Shield I and Shield II phage defense mechanism. **(a)** ShdA normally resides in the membrane, and in absence of a viral challenge, it binds chromosomal DNA without degrading it. Upon injection of phage DNA, the perturbation of the bacterial cytoplasmic membrane results in the activation of ShdA nuclease activity, resulting in chromosomal and phage DNA degradation. **(b)** In the case of Shield II, following membrane perturbation, ShdA II is activated but maintained at low levels by ShdB II. Whilst this can be sufficient to prevent replication of some types of phages, the lower efficiency of ShdA II nuclease activity could result in insufficient DNA degradation and accumulation of specific phage proteins/triggers that can titrate ShdB II, inhibiting its ability to degrade ShdA II, thereby favouring the enrichment of ShdA II and promoting phage DNA and nucleoid degradation and cell death. This figure was created with BioRender.com.

Interestingly, we found a subset of ShdA homologues encoded within a DISARM locus, divergent from Shield-related ShdA proteins (Figure S3). These could potentially represent a fifth additional Shield subtype found associated with DISARM, cooperating with the latter to offer a wider spectrum of protection. A similar cooperation has been observed to some RM systems and other defence systems (2, 44–46).

Recent reports highlighted how anti-phage system can adopt similar protein folds/ domains, in different combinations, to mediate immunity (10, 11, 13, 14). A recent example is represented by the CBASS effector NucC, which was also adopted by a subset of CRISPR-Cas systems (40). Thus, the DISARM-associated ShdA homologues could alternatively represent a case where a DISARM accessory component has acquired a RmuC domain to contribute to phage defense, independently from Shield subtypes. A RmuC-like domain was previously associated with few members of a group of prokaryotic reverse-transcriptases involved in phage defense. However in these cases, the reverse-transcriptase modules were the only ones necessary for defense and the role of RmuC was not explored (47). Finally, the presence of a RmuC-like domain was also predicted for anti-phage systems PD-T7-1 and PD-T7-5, found in *E. coli* (14), but with a much lower probability score and coverage than ShdA II (AN400_RS26690).Furthermore, PD-T7-1 and PD-T7-5 show no sequence similarity to ShdA II, nor a shared predicted fold. Future investigation will establish whether the loci shown in Figure S3 represent another Shield subtype or an example where RmuC domains where independently acquired for bacterial immunity by DISARM components.

Nevertheless, these reports indicate that RmuC domains have been independently acquired for bacterial immunity and our data demonstrate that in Shield systems the RmuC plays a direct and central role in phage inhibition.

To date, despite being functional in *E. coli,* Shield homologues are only found in *Pseudomonas spp.* While it is difficult to speculate on the reason why this system has not been acquired by other species, other homologues with low sequence but high structural similarities might be present in other genomes. We also note that previous studies showed that the number of antiphage systems per strain, as well as the specific types of systems present can deeply vary within *Pseudomonas* strains (27). Rare distribution of some anti-phage systems has been reported previously (11, 13, 27). Taking into consideration that anti-phage systems acquisition comes with a fitness cost for the host bacterium, usually reflected in loss of virulence factors and/or antibiotic resistance (48, 49), for some systems, their sparse distribution could be reflective of a compromise between high fitness cost/quick development of phage countermeasures, and maintaining the system in circulation within the species ‘pan-immune’ system (1). This is particularly relevant for Shield, as its ability to decrease plasmid uptake could generate a fitness disadvantage (e.g. lack of acquisition of pathogenicity islands or antibiotic resistance genes). This is especially important in the context of species like *P. aeruginosa,* where acquisition of pathogenicity genes through horizontal gene transfer frequently takes place (49, 50).

In conclusion, we describe the discovery and defense mechanism of a previously uncharacterised anti-phage system, which utilises RmuC domains for bacterial immunity. Our study describes a unique defense strategy, adding new information to the complex bacterial immunity landscape.

## MATERIAL AND METHODS

### Bacterial strains, plasmids and culture conditions

The strain *P. aeruginosa* NCTC 11442 (equivalent to ATCC 33350) was grown overnight at 37°C for chromosomal DNA extraction. *E. coli* BL21 (DE3) and MG1655 were grown at 37°C on either solid media or liquid culture, shaking at 200 rpm. For liquid growth, Luria broth (LB) was used as the standard medium. For growth on solid media, LB was supplemented with 1.5% (w/v) or with 0.35% (w/v) agar for solid or soft agar, respectively. When required, LB was supplemented with ampicillin (Amp, 50 μg/mL), chloramphenicol (Chl, 25 μg/mL), isopropyl-β-D-thiogalactopyranoside (IPTG, 0.5 mM), L-arabinose (0.2% w/v) or D-glucose (0.2% w/v). Strains and plasmids used in this study are listed in Table S11.

### Measurement of bacterial growth

Overnight cultures were diluted 1:200 in fresh medium and grown until an OD_600nm_ of 0.3. Cultures were infected with phage at an MOI of 1, 0.1 and 0.01 in a final volume of 200 μL and aliquoted into a 96-well plate. The plate was incubated with continuous shaking in a TECAN infinite nano M+ and absorbance at 600 nm was measured every 20 min.

### DNA manipulation and transformation

The chromosomal DNA of *P. aeruginosa* NCTC 11442 was extracted using the GenElute Bacterial Genomic DNA kit (Merck). Plasmid backbones and inserts for cloning were amplified from the purified chromosomal DNA using Q5 High-Fidelity DNA Polymerase (NEB). PCR products and plasmids were purified with Monarch DNA kits (NEB). Overlapping primers for amplification were designed using the NEBuilder assembly tool (https://nebuilderv1.neb.com/) (Table S12). Plasmids and inserts were assembled using NEBuilder HiFi DNA Assembly (NEB), followed by incubation at 50 °C for 20 min. Point mutations or small deletions in pGM34 and pGM71 (Table 1) were performed using the KLD enzyme mix (NEB)

### Phage propagation and lysate preparation

Phage lysates were diluted in phage buffer (10 mM Tris–HCl pH 7.4, 10 mM MgSO4, 0.1% gelatin) and propagated in *E. coli* DH5α. For this purpose, neat lysates or their dilutions were added to 200 μL of *E. coli* DH5α. The mixture was added to 5 mL of soft agar and poured onto LB agar plates. Plates were incubated at 37 °C overnight. Following incubation, the top agar containing confluent phage plaques was scraped off and added to 3 mL of phage buffer and 500 μL of chloroform. Samples were vortexed for 2 minutes and then incubated at 4 °C for 30 min. Samples were subsequently centrifuged at 4000 x *g* for 20 min and the supernatant collected and added to 100 μL of chloroform for storage.

### Efficiency of plating measurement and fold protection calculation

To measure the efficiency of plating (EOP), 10 μL of phage lysate was added to 200 μL of an overnight culture of *E. coli* MG1655 carrying empty pBAD18 or pBAD18 encoding the Shield II system, ShdAII or ShdB II only. Five mL of soft agar was added to each culture and poured onto LB agar plates. As a control strain, plasmid-free MG1655 was used. EOP was measured as the number of PFU mL^-1^ of a test strain divided by the number of PFU/mL of the control strain.

For the calculation of fold protection values shown in Figure 5a serial dilution plaque assays were used. The number of plaques formed on a strain containing the Shield II system was compared to the number of plaques formed on a strain containing the empty vector. The ratio between the two values was used to calculate the fold protection.

### Measurement of phage replication

Measurement of phage replication was performed as previously described (34). Briefly, *E. coli* MG1655 carrying pBAD18 or the same plasmid encoding the Shield II, ShdA only or ShdB only was grown to an OD_600nm_ of 0.6 and infected with 100 PFU/mL of phage ϕSipho or ϕTB34. At time zero and following 90 minutes incubation at 37 °C, chloroform was added to each culture. Cells were then vortexed and centrifuged at 13000 x *g* for 10 min. The supernatant was collected and serially diluted using phage buffer. Subsequently, 10 μL of the chosen dilution was added to 200 μL of the test strain (*E. coli* DH5α) in 5 mL of soft agar and poured onto a LB plate. Plates were incubated overnight at 37 °C and PFU/ mL assessed. Fold phage replication was calculated as: titre (PFU mL^-1^) at 90 min/ titre (PFU mL^-1^) at 0 min.

### Efficiency of centre of infection and cell survival measurements

To measure cell survival after phage infection and the efficiency of centre of infection (ECOI), *E. coli* MG1655 harbouring pBAD18 or the same vector encoding Shield II, ShdA II or ShdB II was grown in LB with ampicillin and 0.2% arabinose to an OD_600nm_ of ~ 0.6 and infected with ϕSipho at MOI of 0.1 for ECOI measurement and MOI 2 for cell survival assessment. Infected cells were incubated for 15 min at 37 °C shaking at 200 rpm to favour adsorption. Following incubation, cells were washed with ice-cold phage buffer twice and then serially diluted in phage buffer. For ECOI, 10 μL of the desired dilution was added to 200 μL of *E. coli* DH5α in 5 mL of soft agar. Infective centres were measured as the number of PFUs on each plate. For cell survival measurement, following serial dilution, 100 μL of the chosen dilution was plated on LB agar and surviving cells were measured as the number of colony forming units (CFU) per plate. The percentage of surviving cells was calculated as: CFU of infected strain/ CFU of non-infected strain.

### One step growth curves

Burst size was calculated as previously reported (5). Briefly, *E. coli* MG1655 carrying pBAD18 or the same vector encoding Shield II, ShdA II or ShdB II was grown to an OD_600nm_ of ~0.6. ϕSipho was added to an MOI of 0.1. Cultures were incubated at 37 °C at 200 rpm and duplicate samples were collected at t = 0 min, t = 30 min and t = 60 min post infection. For each timepoint, one sample was immediately serially diluted and plated onto *E. coli* DH5α top lawns, accounting for free phages and phage-infected cells. The second duplicate sample of each timepoint was instead treated with chloroform before being plated as above, to measure free phages and phage released by infected cells. Burst size was calculated as number of released phages from infected cells (PFU mL^-1^ (t = 60 min) - PFU mL^-1^ (t = 30 min))/ number of phage-infected cells (PFU mL^-1^ (t = 0 min) - PFU mL^-1^ (t = 30 min)).

### Bacterial two hybrid assay

ShdA II and ShdB II were fused to the two fragments of adenylate cyclase encoded on plasmids pT25 and pUT18 (39). Inserts were cloned by the NEBuilder assembly method (NEB). Plasmids were introduced into *E. coli* BTH101 and selected on MacConkey medium (Difco) supplemented with maltose (1%), Amp, 50 μg/mL and Chl 25 μg/mL. Plates were incubated for 48 hrs at 30°C and positive interactions were identified as dark red colonies.

### Cell free protein synthesis and Nuclease assay

The RmuC domains of ShdA I, ShdA II, ShdA III and ShdA IV homologues were synthesised *in vitro* using the PURExpress cell-free transcription/translation kit (NEB), from PCR products. Primers used to generate the appropriate templates are shown in Table S7. Protein synthesis was performed with either 250 ng and 500 ng of DNA template. Synthesis was performed for 4 hr at 37 °C according to the manufacturer’s recommendation. Following incubation, 10 mM MgCl2 was added to the reaction and the final volume was adjusted to 10 μL. Ribosomes were removed through centrifugation for 60 min at 15,000 rpm at 4 °C, through an Amicon Ultracel 0.5mL spin concentrator with a 100 KDa filter (Merck). The flowthrough was collected and the His-tagged PURExpress kit components were removed from the reaction following incubation with Ni-NTA agarose beads (Thermo) for 45 min at 4 °C. Agarose beads were removed through centrifugation with Biorad micro Bio-spin columns at 15,000 g for 10 min at 4 °C. As a control, the same reactions were performed in parallel with the dihydrofolate reductase (DHFR) control provided by the PURExpress kit.

To test for nuclease activity the in vitro synthesised ShdA proteins or DHFR control were incubated with 20 ng of phage ϕSipho, *E. coli* MG1655 chromosome or plasmid pSG483 DNA, followed by agarose gel electrophoresis and staining with GelRed (Cambridge Bioscience).

### Fluorescence microscopy and quantification of DAPI fluorescence

Overnight cultures (5 ml) were diluted into 25 mL LB containing 0.2% arabinose and 50 μg/mL ampicillin and grown for 2 hr. 200 μL of each culture were collected at timepoints t=0 hr, t=1 hr and t=2 hr and stained with 4’,6-diamidino-2-phenylindole (DAPI) at a final concentration of 5 μg /mL and F M5-95(Thermo) at 200 μg /mL. Staining was carried out at 37 °C for 15 min and then 2 μL of each culture was transferred on a microscope slide with a pad of 1 % UltraPure agarose (Invitrogen) in H2O. Imaging was carried out on Nikon Eclipse Ti equipped with CoolLED pE-300^white^ light source, Nikon Plan Apo 100×/1.40 NA Oil Ph3 objective, and Photometrics Prime sCMOS, and Chroma 49008 (Ex 560/40, Dm 585, Em 630/75) filter set for FM 5-95 and Chroma 49000 (Ex 350/50, DM 400, EM 460/50) filter set for DAPI. The images were captured using Metamorph 7.7 (Molecular Devices) and analysed using Fiji (51).

Quantification of DAPI fluorescence was performed using Fiji. In brief, individual cells were identified from thresholded phase contrast images and converted to regions of interest (ROI). The ROI area associated with the cell periphery was defined as 3 pixel wide band extending towards the cell interior. These whole cell and cell periphery -ROIs, and background-subtracted fluorescence images were then used quantify the integrated density of DAPI fluorescence signal for the whole cell, and for the respective cell periphery. At last, a ratio of the fluorescence signals between the cell periphery and the whole cell was calculated and depicted in a swarm plot.

### ShdA II detection

ShdA II with a C-terminal His_6_ tag was expressed from the arabinose inducible plasmid, pBAD18 alone or in presence of ShdB II. Induction was performed with 0.2% L-arabinose for 5 hrs. Following induction, cells were harvested by centrifugation at 14,000 x g for 10 min at 4 °C and resuspended in Laemmli buffer. His_6_ tagged proteins were detected with anti-His_6_ primary antibody (Pierce; 1:6,000) and anti-mouse secondary antibody (Biorad; 1:10,000). GroEL was detected with anti-GroEL primary antibody (Pierce; 1:10,000) and anti-rabbit secondary antibody (Biorad; 1:20,000).

### Subcellular fractionation

Overnight cultures of MG1655 carrying ShdA II with a C-terminal His6 tag were diluted in 25 mL LB containing 0.2 % L-arabinose and grown for 5 hrs. Following growth, cells were centrifuged at 4000 x g for 10 min at 4 °C. Cells were resuspended in 1 mL of Tris HCl pH 8 and lysed by sonication. Cellular debris were removed by centrifugation at 14,000 x g for 10 min at 4 °C. The supernatant was collected and subjected to ultracentrifugation (200,000 x g, 30 min, 4 °C). The supernatant was collected, representing the cytoplasmic fraction and the pellet resuspended in 1 mL of Tris HCl pH 8, representing the total membrane fraction. His6 tagged proteins were detected with anti-His6 primary antibody (Pierce; 1:6,000) and antimouse secondary antibody (Biorad; 1:10,000). GroEL was used as cytoplasmic control and detected as above. TatA was used as membrane control protein and detected with anti-TatA serum as previously described (52).

### Homology searches and gene neighbourhood analysis

ShdA alignments were generated using MUSCLE (v3.8.1551) (53). Alignments were used to generate Hidden Markov Models with the HMMER suite (v 3.3.2) (54). The obtained models were used to query a local Refseq database of *Pseudomonas spp* or all bacterial genomes.The cutoff value was set to a bit score of 30 over the overall sequence/profile comparison. Efetch from the entrez utilities (55) was employed to retrieve an identical protein group report (IPG) for each hit protein obtained from the HMMER searches. Neighboring genes to Shield system were identified using FlaGs version 1.2.7 (29) with a non-redundant ShdA protein set as query. Neighboring genes were analysed using the defense finder tool to determine their association to known anti-phage systems.

### Annotation of anti-phage systems

Flanking genes of Shield systems were retrieved using the FlaGs tool (29). Clustered flanking genes retrieved by FlaGs were subjected to searches against a local PFAM database (56). Additionally, assembly IDs of genomes encoding Shield homologues, retrieved using Efetch, were used to download genome proteomes. Proteomes were then used as an input to predict anti-phage systems with defense finder (27). Genome neighbourhoods were scanned and anti-phage system predictions were manually curated by comparing defense finder results with PFAM predictions (56). For multi-gene loci where PFAM and defense finder predictions were discordant, the prediction that best matched their operon organisation was chosen.

### Protein function prediction

Where possible, protein function predictions were performed using a local PFAM database (56). Protein structures were predicted using Alphafold and the Dali server (50, 51). Presence of signal peptides and transmembrane domains were predicted using DeepTMHMM and SignalP 6.0 (57, 58).

### Phylogenetic analysis

The alignment of ShdA proteins was used to build a maximum likelihood phylogenetic tree with IQTREE (v 2.1.4) (59) with 1000 ultrafast bootstraps. Trees were plotted and annotated in iTOL (60).

## Author contributions

G.M., T.P. and T.R.B conceived the study. G.M., T.R.B and H.S. designed experiments and analysed the data. G.M and E.M performed experimental work and bioinformatic analyses. G.M., T.P., and T.R.B. wrote the paper with inputs from all other authors.

## Acknowledgements

We are grateful to Newcastle University Molecular Microbiology MRes 4828F programme for supporting E.M. We thank Stephen Garrett and Eleanor Boardman for helpful advice and discussion. We are grateful to Dr James Connolly for the gift of anti-GroEL antibody.

