## Supplementary Figures and Tables S11-12 for "Shield co-opts an RmuC domain to mediate phage defence across *Pseudomonas* species"

PNAS

**Supporting Information for**
Shield co-opts an RmuC domain to mediate phage defence across Pseudomonas species.

Elliot Macdonald, Henrik Strahl, Tim R. Blower, Tracy Palmer and Giuseppina Mariano
Giuseppina Mariano*

*to whom correspondence should be addressed:
Giuseppina Mariano,

**This PDF file includes:**

Figures S1 to S7
Tables S11 to S12
SI References

**Other supporting materials for this manuscript include the following:**

Tables S1 to S10 (Provided separately)

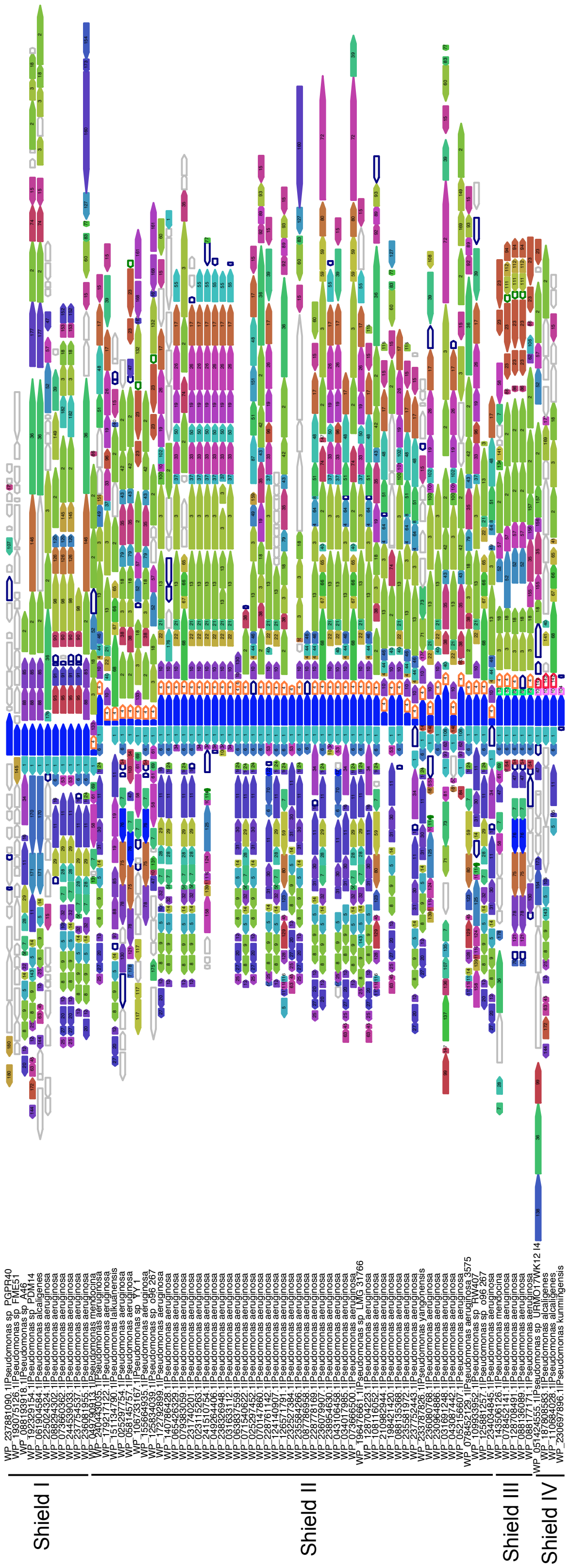


**Figure S1. Definition of Shield subtypes.** FlaGs output representing the genomic neighbourhood of *shdA* genes belonging to different subtypes. The spectrum of genomes encoding these homologues is reported in Tables S2-S5. FlaGs-grouped genes are numbered and coloured by FlaGs according to their association to a certain cluster. FlaGs-numbering of clustered genes is reported in Table S4. Clustered genes were annotated as belonging to a specific antiphage system using PFAM and defense-finder. These annotations are reported in Table S4. *shdA* genes are coloured in blue and partner genes have a coloured outline as indicated on the figure.

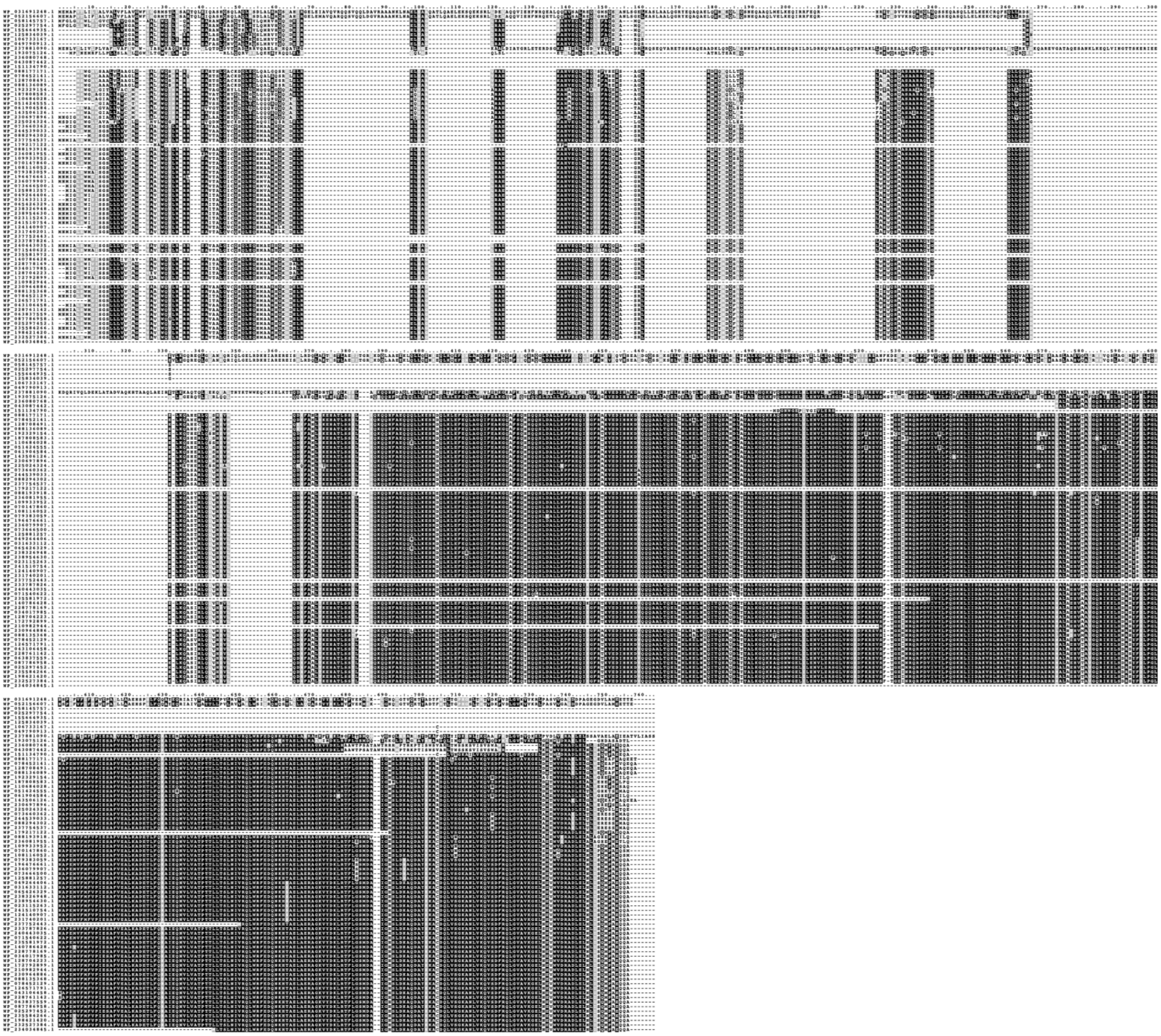

**Figure S2. Alignment of ShdA homologues from different subtypes.** Alignment of ShdA homologues using MUSCLE. The alignment was visualised in Boxshade and coloured by percentage of identity.

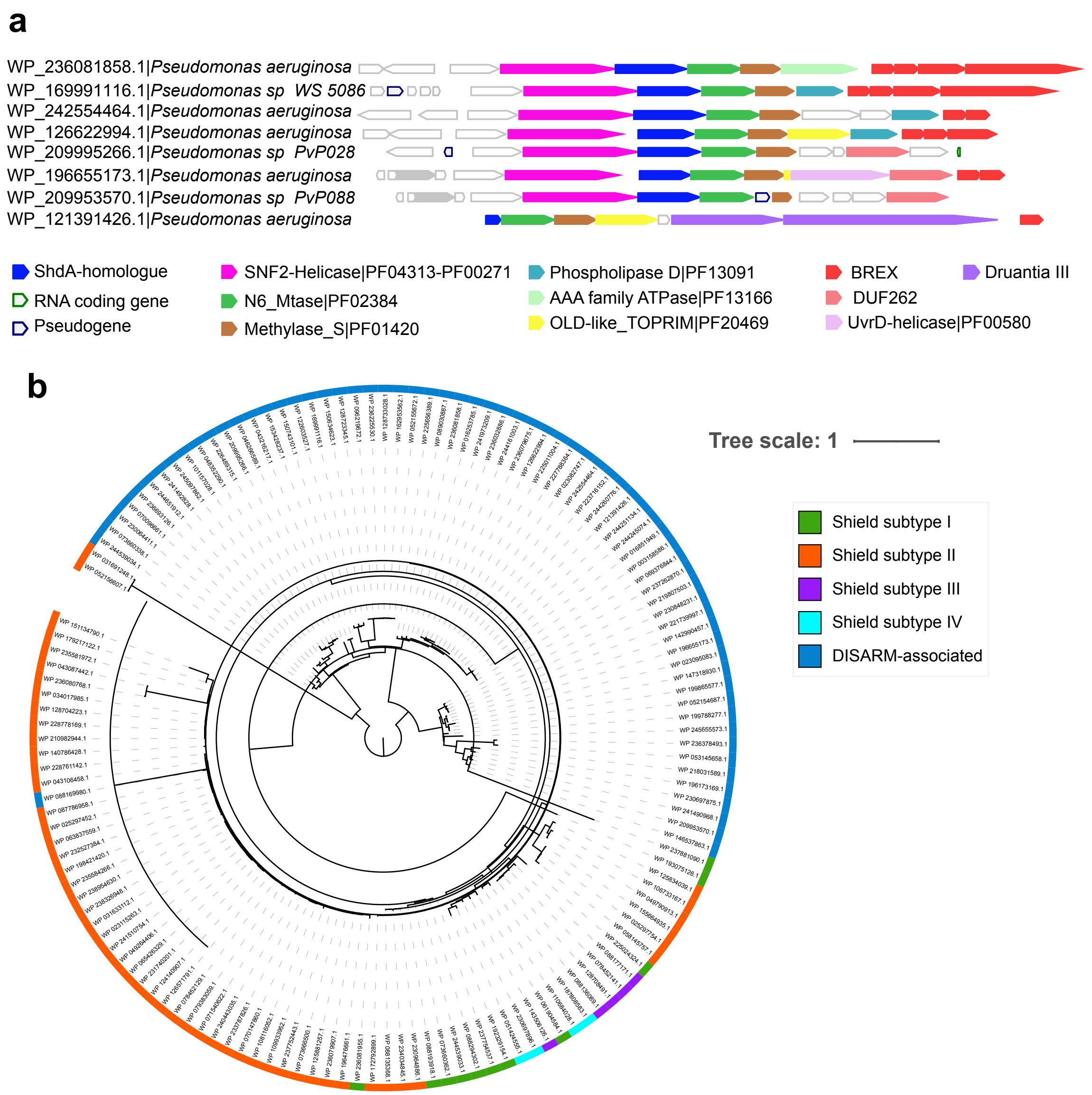


**Figure S3: ShdA homologues are encoded in DISARM loci. (a)** Schematic representation of ShdA homologues encoded within DISARM operons. The full set of DISARM-associated ShdA loci is shown in Figure S4. **(b)** Phylogenetic tree based on the ShdA homologues from Figure S1 in addition to DISARM-associated ShdA. Coloured blocks were used to show ShdA homologues belonging to distinct Shield subtypes or DISARM-like loci.

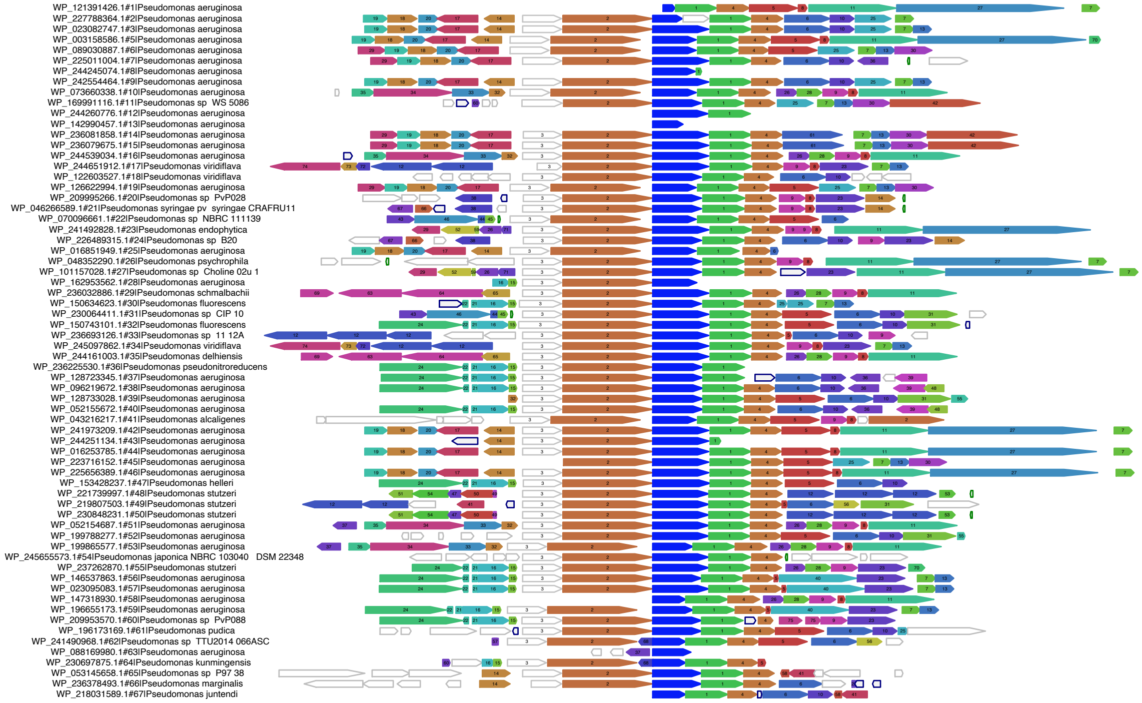


**Figure S4. Distant ShdA homologues can be encoded in DISARM-like operons.** FlaGs output representing the genomic neighbourhood of distant ShdA homologues shows that these can be encoded within DISARM-like operons. Neighbouring genes are clustered together based on similarity and each cluster is numbered. Definition of neighbouring gene clusters is reported in Table S8

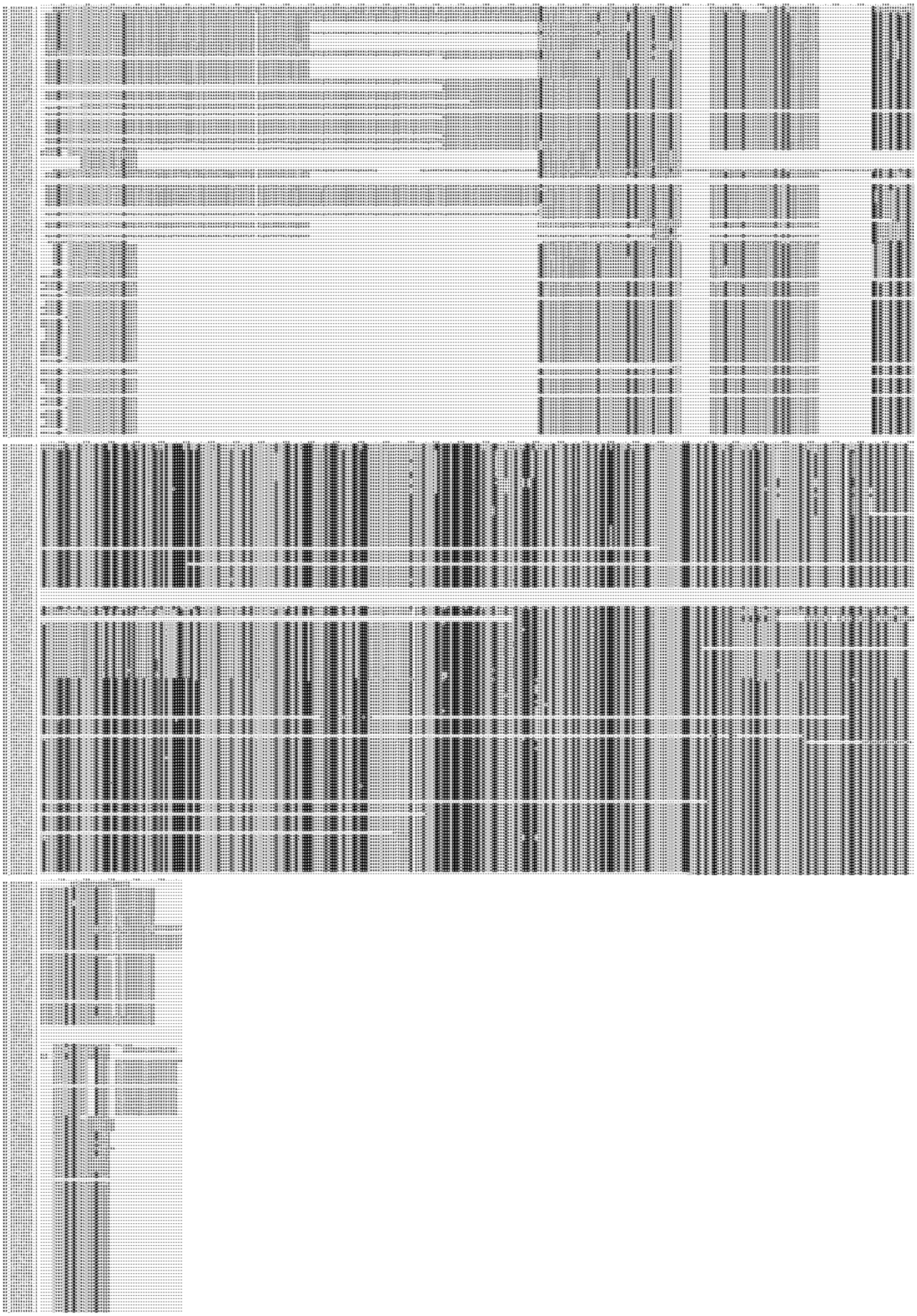

**Figure S5: Alignment of ShdA homologues from different Shield subtypes and DISARM-associated ShdA.** Alignment of ShdA II homologues using MUSCLE. The alignment was coloured in Boxshade by percentage of identity. Sequence conservation between Shield- and DISARM-associated ShdA homologues is still present but is reduced to what observed in Figure S2.

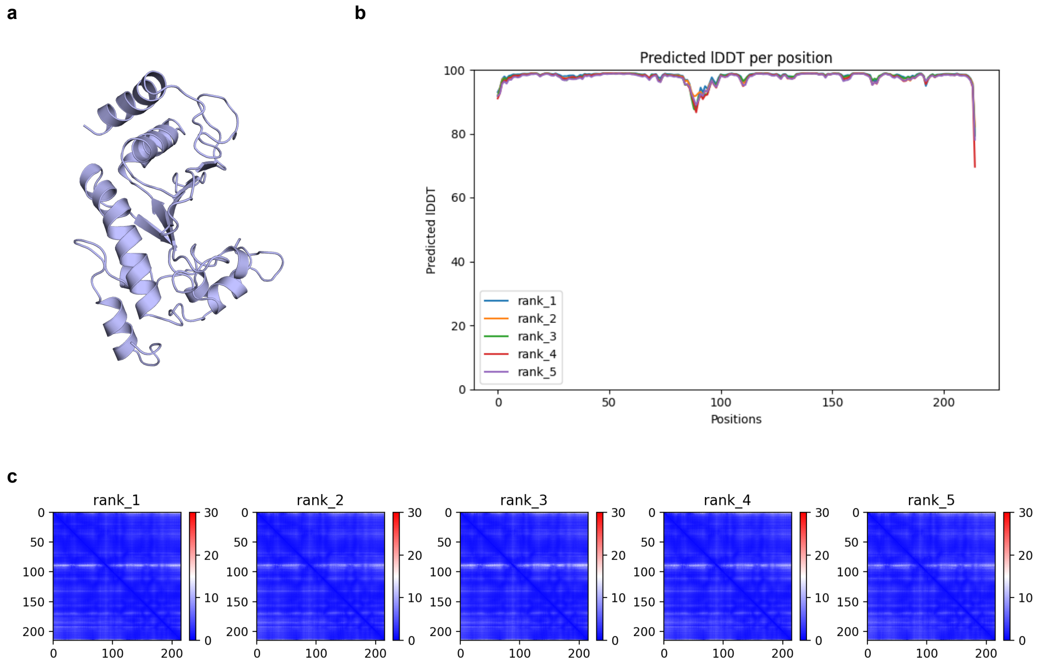


**Figure S6. ShdB II structural prediction suggests a peptidase M15 fold. (a)** Alphafold predicted structure of ShdB II was used for Dali predictions (Table S4), showing it harbours a predicted peptidase M15 domain. **(b)** Local Distance Difference Test (lDDT) relative to ShdB II predicted structure. lDDT shows a per-residue measure of local confidence for the prediction and it is high across the whole ShdB II structure. **(c)** Predicted Aligned Error (PAE) for ShdB II Alphafold-predicted structure. PAE reports the expected error at each residue position. For ShdB II structure PAE was low across the whole sequence.

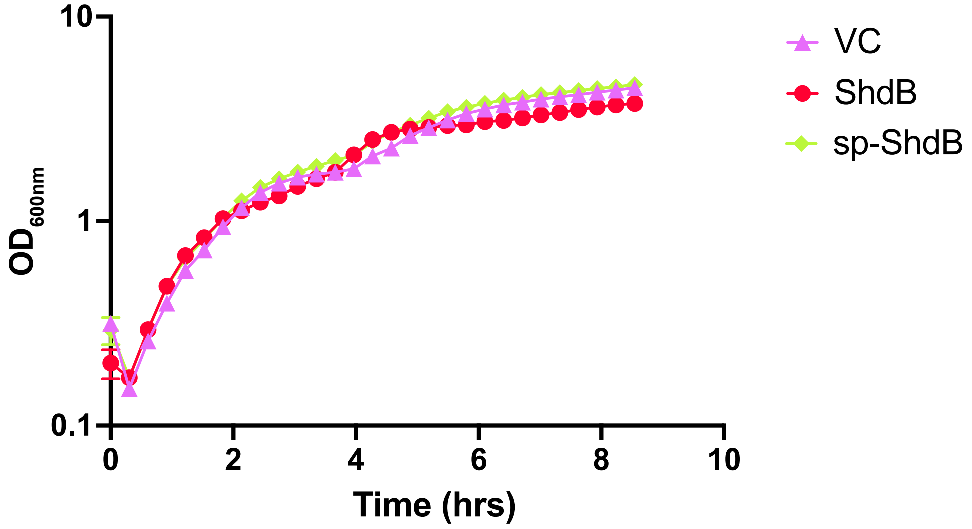


**Figure S7: ShdB II expression with a signal peptide does not impair cell growth.** Growth in liquid LB media of *E. coli* MG1655 carrying (VC, pBAD18), ShdB II or ShdB II with an OmpA signal sequence (sp-ShdB II). Expression was repressed with 0.2% D-glucose or induced with addition of 0.2% L-arabinose. Points show mean +/− SEM (n = 3 biological replicates).

**Table S11.** Strains and plasmids used in this study

| Name | Description | Reference |
|---|---|---|
| **Strains** | | |
| ***Pseudomonas aeruginosa*** | *Pseudomonas aeruginosa* strain NCTC 11442 | NCTC |
| ***Escherichia coli*** | | |
| MG1655 | Wild type (model K-12 strain) | (1) |
| DH5α | Cloning strain, F– φ80lacZΔM15 Δ(lacZYA-argF)U169 recA1 endA1 hsdR17(rK–, mK+) phoA supE44 λ–thi-1 gyrA96 relA1 | New England Biolabs |
| **Plasmids** | | |
| pBAD18 | Arabinose-inducible expression vector ($Kn^R$); gene of interest is cloned downstream of the $P_{ara}$ promoter | (2) |
| pUT18 | Bacterial Two Hybrid plasmid (for fusion of target protein with C-terminal T18 fragment of CyaA; $Amp^R$) | (3) |
| pT25 | Bacterial Two Hybrid plasmid (for fusion of target protein with Nterminal T25 fragment of CyaA; $Cm^R$) | (3) |
| pGM34 | Coding sequence for Shield II (AN400_RS26690 and AN400_RS26695) from *P. aeruginosa* ATCC33350 in pBAD18 | This study |
| pGM42 | Coding sequence for ShdB II (AN400_RS26695) from *P. aeruginosa* ATCC33350 in pBAD18 | This study |

| | | |
|---|---|---|
| pGM43 | Coding sequence for ShdA II (AN400_RS26690) from *P. aeruginosa* ATCC33350 in pBAD18 | This study |
| pGM122 | Coding sequence for ShdA I (DL351_RS13220) from *P. aeruginosa* BH9 in pBAD18 | This study |
| pGM133 | Coding sequence for Shield III (EQ826_RS12590, EQ826_RS12595 and EQ826_12600) from *P. mendocina* FFL34 in pBAD18 | This study |
| pGM134 | Coding sequence for ShdA III (EQ826_RS12590) from *P. mendocina* FFL34 in pBAD18 | This study |
| pGM126 | Coding sequence for Shield IV (A9179_RS12860, A9179_RS12855, A9179_RS12850) from *P. alcaligenes* AVO110 in pBAD18 | This study |
| pGM127 | Coding sequence for ShdA IV (A9179_RS12860) from *P. alcaligenes* AVO110 in pBAD18 | This study |
| pGM128 | Coding sequence for ShdD (A9179_RS12855) from *P. alcaligenes* AVO110 in pBAD18 | This study |
| pGM139 | Coding sequence for ShdE (A9179_RS12850) from *P. alcaligenes* AVO110 in pBAD18 | This study |
| pGM130 | Coding sequence for ShdAD (A9179_RS12860, A9179_RS12855) from *P. alcaligenes* AVO110 in pBAD18 | This study |
| pGM131 | Coding sequence for ShdAE(A9179_RS12860, A9179_RS12850) from *P. alcaligenes* AVO110 in pBAD18 | This study |

| | | |
|---|---|---|
| pGM132 | Coding sequence for ShdDE (A9179_RS12855, A9179_RS12850) from *P. alcaligenes* AVO110 in pBAD18 | This study |
| pGM107 | Coding sequence for ShdA II (AN400_RS26690) fused with a C-terminal $His_6$ tag and coding sequence of ShdB II (AN400_RS26695) in pBAD18 | This study |
| pGM116 | Coding sequence for ShdA II (AN400_RS26690) fused with a C-terminal $His_6$ tag in pBAD18 | This study |
| pGM117 | Coding sequence for ShdA II (AN400_RS26690) in pUT18 | This study |
| pGM118 | Coding sequence for ShdB II (AN400_RS26695) in pUT18 | This study |
| pGM119 | Coding sequence for ShdA II (AN400_RS26690) in pT25 | This study |
| pGM120 | Coding sequence for ShdB II (AN400_RS26695) in pT25 | This study |
| pUT18-NarG | Coding sequence of NarG in pUT18 | (4) |
| pT25-NarJ | Coding sequence of NarJ in pT25 | (4) |

**Table S12.** Oligonucleotide primers and additional details for plasmid construction.

| Plasmid | Sequence of relevant primers (5'-3')[a] | Description |
|---|---|---|
| pGM34 | GGTACCCGGGGATCCTC | Forward primer to clone AN400_RS26690 and AN400_RS26695 in pBAD18 by NEBuilder HiFi DNA Assembly. Primer is used to linearise vector |
| | GAATTCGCTAGCCCAAAAAAAC | Reverse primer to clone AN400_RS26690 and AN400_RS26695 in pBAD18 by NEBuilder HiFi DNA Assembly. Primer is used to linearise vector |
| | GTTTTTTTGGGCTAGCGAATTCATGATTGGATTGTCGTGGATAGTGATTAG | Forward primer to clone AN400_RS26690 and AN400_RS26695 in pBAD18 by NEBuilder HiFi DNA Assembly. Primer is used to amplify genes of interest |
| | AGGATCCCCGGGTACCCTAGTTGAGACTCGCCAACCATGGGC | Reverse primer to clone AN400_RS26690 and AN400_RS26695 in pBAD18 by NEBuilder HiFi DNA Assembly. Primer is used to amplify genes of interest |
| pGM43 | GGTACCCGGGGATCCTC | Forward primer to clone AN400_RS26690 in pBAD18 by NEBuilder HiFi DNA Assembly. Primer is used to linearise vector |
| | GAATTCGCTAGCCCAAAAAAAC | Reverse primer to clone AN400_RS26690 in pBAD18 by NEBuilder HiFi DNA Assembly. Primer is used to linearise vector |
| | GTTTTTTTGGGCTAGCGAATTCATGATTGGATTGTCGTGGATAGTGATTAG | Forward primer to clone AN400_RS26690 in pBAD18 by NEBuilder HiFi DNA Assembly. Primer is used to amplify genes of interest |
| | AGGATCCCCGGGTACCTCACGCCTGCTGCTCAG | Reverse primer to clone AN400_RS26690 in pBAD18 by NEBuilder HiFi DNA Assembly. Primer is used to amplify genes of interest |
| pGM42 | GGTACCCGGGGATCCTC | Forward primer to clone AN400_RS26695 in pBAD18 by NEBuilder HiFi DNA Assembly. Primer is used to linearise vector |
| | GAATTCGCTAGCCCAAAAAAAC | Reverse primer to clone AN400_RS26695 in pBAD18 by NEBuilder HiFi DNA Assembly. Primer is used to linearise vector |

| | | |
|---|---|---|
| | CGTTTTTTTGGGCTAGCGAATTCATGAAAAAACCAGCAACGGTAC | Forward primer to clone AN400_RS26690 in pBAD18 by NEBuilder HiFi DNA Assembly. Primer is used to amplify genes of interest |
| | AGGATCCCCGGGTACCTCACGCCTGCTGCTCAG | Reverse primer to clone AN400_RS26695 in pBAD18 by NEBuilder HiFi DNA Assembly. Primer is used to amplify genes of interest |
| pGM122* | GAATTCGCTAGCCCAAAAAAAC | Forward primer to clone DL351_RS13220 pBAD18 by NEBuilder HiFi DNA Assembly. Primer is used to linearise vector |
| | GGTACCCGGGGATCCTCTAG | Reverse primer to clone DL351_RS13220 in pBAD18 by NEBuilder HiFi DNA Assembly. Primer is used to linearise vector |
| | CGTTTTTTTGGGCTAGCGAATTCATGTCCTGGGCAGTAATG | Forward primer to clone DL351_RS13220 in pBAD18 by NEBuilder HiFi DNA Assembly. Primer is used to amplify genes of interest |
| | CTAGAGGATCCCCGGGTACCTTACTGCGCCATTTCGCTG | Reverse primer to clone DL351_RS13220 in pBAD18 by NEBuilder HiFi DNA Assembly. Primer is used to amplify genes of interest |
| pGM133* | GAATTCGCTAGCCCAAAAAAAC | Forward primer to clone EQ826_RS12590, EQ826_RS12595 and EQ826_12600 in pBAD18 by NEBuilder HiFi DNA Assembly. Primer is used to linearise vector |
| | GGTACCCGGGGATCCTCTAG | Reverse primer to clone EQ826_RS12590, EQ826_RS12595 and EQ826_12600 in pBAD18 by NEBuilder HiFi DNA Assembly. Primer is used to linearise vector |
| | CGTTTTTTTGGGCTAGCGAATTCATGTCCTGGGCAGTAATG | Forward primer to clone EQ826_RS12590, EQ826_RS12595 and EQ826_12600 in pBAD18 by NEBuilder HiFi DNA Assembly. Primer is used to amplify genes of interest |
| | CTAGAGGATCCCCGGGTACCTCAGCCCAGGCTAGCTAG | Reverse primer to clone EQ826_RS12590, EQ826_RS12595 and EQ826_12600 in pBAD18 by NEBuilder HiFi DNA Assembly. Primer is used to amplify genes of interest |
| pGM134* | GAATTCGCTAGCCCAAAAAAAC | Forward primer to clone EQ826_RS12590 in pBAD18 by NEBuilder HiFi DNA Assembly. Primer is used to linearise vector |
| | GGTACCCGGGGATCCTCTAG | Reverse primer to clone EQ826_RS12590 in pBAD18 by |

| | | |
|---|---|---|
| | | NEBuilder HiFi DNA Assembly. Primer is used to linearise vector |
| | CGTTTTTTTGGGCTAGCGAATTCATGTCCTGGGCAGTAATGGG | Forward primer to clone EQ826_RS12590in pBAD18 by NEBuilder HiFi DNA Assembly. Primer is used to amplify genes of interest |
| | CTAGAGGATCCCCGGGTACCTGCTTCCTCCTGCGCCTC | Reverse primer to clone EQ826_RS12590in pBAD18 by NEBuilder HiFi DNA Assembly. Primer is used to amplify genes of interest |
| pGM126* | GAATTCGCTAGCCCAAAAAAAC | Forward primer to clone A9179_RS12860, A9179_RS12855 and A9179_RS12850 in pBAD18 by NEBuilder HiFi DNA Assembly. Primer is used to linearise vector |
| | GGTACCCGGGGATCCTCTAG | Reverse primer to clone A9179_RS12860, A9179_RS12855 and A9179_RS12850 in pBAD18 by NEBuilder HiFi DNA Assembly. Primer is used to linearise vector |
| | CGTTTTTTTGGGCTAGCGAATTCATGTCTTGGGGAGTAGCGG | Forward primer to clone A9179_RS12860, A9179_RS12855 and A9179_RS12850 in pBAD18 by NEBuilder HiFi DNA Assembly. Primer is used to amplify genes of interest |
| | CTAGAGGATCCCCGGGTACCTCATCCTTGGCCGGCGAG | Reverse primer to clone A9179_RS12860, A9179_RS12855 and A9179_RS12850 in pBAD18 by NEBuilder HiFi DNA Assembly. Primer is used to amplify genes of interest |
| pGM127* | GAATTCGCTAGCCCAAAAAAAC | Forward primer to clone A9179_RS12860 in pBAD18 by NEBuilder HiFi DNA Assembly. Primer is used to linearise vector |
| | GGTACCCGGGGATCCTCTAG | Reverse primer to clone A9179_RS12860 in pBAD18 by NEBuilder HiFi DNA Assembly. Primer is used to linearise vector |
| | CGTTTTTTTGGGCTAGCGAATTCATGTCTTGGGGAGTAGCGG | Forward primer to clone A9179_RS12860 in pBAD18 by NEBuilder HiFi DNA Assembly. Primer is used to amplify genes of interest |
| | CTAGAGGATCCCCGGGTACCTTACTGCGGCAATTCGCTG | Reverse primer to clone A9179_RS12860in pBAD18 by NEBuilder HiFi DNA Assembly. Primer is used to amplify genes of interest |
| pGM128* | GAATTCGCTAGCCCAAAAAAAC | Forward primer to A9179_RS12855 in pBAD18 by NEBuilder HiFi DNA Assembly. Primer is used to linearise vector |

| | | |
|---|---|---|
| | GGTACCCGGGGATCCTCTAG | Reverse primer to clone A9179_RS12855 in pBAD18 by NEBuilder HiFi DNA Assembly. Primer is used to linearise vector |
| | CGTTTTTTTGGGCTAGCGAATTCATGCAGGAGTCGTACGACTTTG | Forward primer to clone A9179_RS12855 in pBAD18 by NEBuilder HiFi DNA Assembly. Primer is used to amplify genes of interest |
| | CTAGAGGATCCCCGGGTACCCTACCAACCTCGACGGGC | Reverse primer to clone A9179_RS12855 in pBAD18 by NEBuilder HiFi DNA Assembly. Primer is used to amplify genes of interest |
| pGM129 * | GAATTCGCTAGCCCAAAAAAAC | Forward primer to clone A9179_RS12850 in pBAD18 by NEBuilder HiFi DNA Assembly. Primer is used to linearise vector |
| | GGTACCCGGGGATCCTCTAG | Reverse primer to clone A9179_RS12850 in pBAD18 by NEBuilder HiFi DNA Assembly. Primer is used to linearise vector |
| | CGTTTTTTTGGGCTAGCGAATTCATGCAGTCGTTGCATATAAGACAGC | Forward primer to clone A9179_RS12850 in pBAD18 by NEBuilder HiFi DNA Assembly. Primer is used to amplify genes of interest |
| | CTAGAGGATCCCCGGGTACCTCATCCTTGGCCGGCGAG | Reverse primer to clone A9179_RS12850 in pBAD18 by NEBuilder HiFi DNA Assembly. Primer is used to amplify genes of interest |
| pGM130 * | GAATTCGCTAGCCCAAAAAAAC | Forward primer to clone A9179_RS12860 and A9179_RS12855 in pBAD18 by NEBuilder HiFi DNA Assembly. Primer is used to linearise vector |
| | GGTACCCGGGGATCCTCTAG | Reverse primer to clone A9179_RS12860 and A9179_RS12855 in pBAD18 by NEBuilder HiFi DNA Assembly. Primer is used to linearise vector |
| | CGTTTTTTTGGGCTAGCGAATTCATGTCTTGGGGAGTAGCGG | Forward primer to clone A9179_RS12860 and A9179_RS12855 in pBAD18 by NEBuilder HiFi DNA Assembly. Primer is used to amplify genes of interest |
| | CTAGAGGATCCCCGGGTACCCTACCAACCTCGACGGGC | Reverse primer to clone A9179_RS12860 and A9179_RS12855 in pBAD18 by NEBuilder HiFi DNA Assembly. Primer is used to amplify genes of interest |
| pGM131 * | GAATTCGCTAGCCCAAAAAAAC | Forward primer to clone A9179_RS12860 and A9179_RS12850 in pBAD18 by NEBuilder HiFi DNA Assembly. Primer is used to linearise vector |
| | GGTACCCGGGGATCCTCTAG | Reverse primer to clone A9179_RS12860 and A9179_RS12850 in pBAD18 by |

| | | |
|---|---|---|
| | | NEBuilder HiFi DNA Assembly. Primer is used to linearise vector |
| | CGTTTTTTTGGGCTAGCGAATTCATGTCTTGGGGAGTAGCGG | Forward primer to clone A9179_RS12860 and A9179_RS12850 in pBAD18 by NEBuilder HiFi DNA Assembly. Primer is used to amplify A9179_RS12860 |
| | ACGACTGCATTTACTGCGGCAATTCGCTG | Reverse primer to A9179_RS12860 and A9179_RS12850 in pBAD18 by NEBuilder HiFi DNA Assembly. Primer is used to amplify A9179_RS12860 |
| | GCCGCAGTAAATGCAGTCGTTGCATATAAGACAGC | Forward primer to A9179_RS12860 and A9179_RS12850 in pBAD18 by NEBuilder HiFi DNA Assembly. Primer is used to amplify A9179_RS12850 |
| | CTAGAGGATCCCCGGGTACCTCATCCTTGGCCGGCGAG | Reverse primer to A9179_RS12860 and A9179_RS12850 in pBAD18 by NEBuilder HiFi DNA Assembly. Primer is used to amplify A9179_RS12850 |
| pGM132* | GAATTCGCTAGCCCAAAAAAAC | Forward primer to clone A9179_RS12855 and A9179_RS12850 in pBAD18 by NEBuilder HiFi DNA Assembly. Primer is used to linearise vector |
| | GGTACCCGGGGATCCTCTAG | Reverse primer to clone A9179_RS12855 and A9179_RS12850 in pBAD18 by NEBuilder HiFi DNA Assembly. Primer is used to linearise vector |
| | CGTTTTTTTGGGCTAGCGAATTCATGCAGGAGTCGTACGACTTTG | Forward primer to clone A9179_RS12855 and A9179_RS12850 in pBAD18 by NEBuilder HiFi DNA Assembly. Primer is used to amplify genes of interest |
| | CTAGAGGATCCCCGGGTACCTCATCCTTGGCCGGCGAG | Reverse primer to clone A9179_RS12855 and A9179_RS12850 in pBAD18 by NEBuilder HiFi DNA Assembly. Primer is used to amplify genes of interest |
| pGM117 | AGTCGACCTGCAGGCATG | Forward primer to clone AN400_RS26690 in pUT18 by NEBuilder HiFi DNA Assembly. Primer is used to linearise vector |
| | CTAGAGGATCCCCGGGTAC | Reverse primer to clone AN400_RS26690 in pUT18 by NEBuilder HiFi DNA Assembly. Primer is used to linearise vector |
| | TGCATGCCTGCAGGTCGACTATGATTGGATTGTCGTGGATAGTGATTAGC | Forward primer to clone AN400_RS26690 in pUT18 by NEBuilder HiFi DNA Assembly. Primer is used to amplify gene of interest |
| | GGTACCCGGGGATCCTCTAGCGCCTGCTGCTCAGCTGG | Reverse primer to clone AN400_RS26690 in pUT18 by NEBuilder HiFi DNA Assembly. Primer is used to amplify genes of interest |

| | | |
|---|---|---|
| pGM118 | AGTCGACCTGCAGGCATG | Forward primer to clone AN400_RS26695 in pUT18 by NEBuilder HiFi DNA Assembly. Primer is used to linearise vector |
| | CTAGAGGATCCCCGGGTAC | Reverse primer to clone AN400_RS26695 in pUT18 by NEBuilder HiFi DNA Assembly. Primer is used to linearise vector |
| | TGCATGCCTGCAGGTCGACTATGAAAAAC CAGCAACGGTAC | Forward primer to clone AN400_RS26695 in pUT18 by NEBuilder HiFi DNA Assembly. Primer is used to amplify gene of interest |
| | GGTACCCGGGGATCCTCTAGGTTGAGACT CGCCAACCATG | Reverse primer to clone AN400_RS26695 in pUT18 by NEBuilder HiFi DNA Assembly. Primer is used to amplify genes of interest |
| pGM119 | AGTCGACCCTGCAGCCCG | Forward primer to clone AN400_RS26690 in pT25 by NEBuilder HiFi DNA Assembly. Primer is used to linearise vector |
| | CTAGAGGATCCCCGGGTAC | Reverse primer to clone AN400_RS26690 in pT25 by NEBuilder HiFi DNA Assembly. Primer is used to linearise vector |
| | GGCGGGCTGCAGGGTCGACTATTGGATTG TCGTGGATAGTGATTAGC | Forward primer to clone AN400_RS26690 in pT25 by NEBuilder HiFi DNA Assembly. Primer is used to amplify gene of interest |
| | GGTACCCGGGGATCCTCTAGTCACGCCTG CTGCTCAGC | Reverse primer to clone AN400_RS26690 in pT25 by NEBuilder HiFi DNA Assembly. Primer is used to amplify genes of interest |
| pGM120 | AGTCGACCCTGCAGCCCG | Forward primer to clone AN400_RS26695 in pT25 by NEBuilder HiFi DNA Assembly. Primer is used to linearise vector |
| | CTAGAGGATCCCCGGGTAC | Reverse primer to clone AN400_RS26695 in pT25 by NEBuilder HiFi DNA Assembly. Primer is used to linearise vector |
| | GGCGGGCTGCAGGGTCGACTAAAAAACCA GCAACG | Forward primer to clone AN400_RS26695 in pT25 by NEBuilder HiFi DNA Assembly. Primer is used to amplify gene of interest |
| | GGTACCCGGGGATCCTCTAGCTAGTTGAGA CTCGCCAAC | Reverse primer to clone AN400_RS26695 in pT25 by NEBuilder HiFi DNA Assembly. Primer is used to amplify genes of interest |
| pGM107 | AGTACTGCCAAGGAGCAAAAC | Forward primer to clone AN400_RS26690 with a C-termina $His_6$ |

| | | |
|---|---|---|
| | | tag in pGM42 upstream of AN400_RS26695 by NEBuilder HiFi DNA Assembly. Primer is used to linearise vector |
| | AGTACTGCCAAGGAGCAAAAC | Reverse primer to clone AN400_RS26690 with a C-termina $His_6$ tag in pGM42 upstream of AN400_RS26695 by NEBuilder HiFi DNA Assembly. Primer is used to linearise vector |
| | CGTTTTTTTGGGCTAGCGAATTCATGATTGGATTGTCGTGG | Forward primer to clone AN400_RS26690 with a C-termina $His_6$ tag in pGM42 upstream of AN400_RS26695 by NEBuilder HiFi DNA Assembly. Primer is used to linearise vector |
| | CTGTTTTGCTCCTTGGCAGTACTTTAGTGATGGTGATGGTG | Reverse primer to clone AN400_RS26690 with a C-termina $His_6$ tag in pGM42 upstream of AN400_RS26695 by NEBuilder HiFi DNA Assembly. Primer is used to linearise vector |
| pGM116 | AGTACTGCCAAGGAGCAAAAC | Forward primer to clone AN400_RS26690 with a C-termina $His_6$ tag in pBAD18 by NEBuilder HiFi DNA Assembly. Primer is used to linearise vector |
| | AGTACTGCCAAGGAGCAAAAC | Forward primer to clone AN400_RS26690 with a C-termina $His_6$ tag in pBAD18 by NEBuilder HiFi DNA Assembly. Primer is used to linearise vector |
| | CCGTTTTTTTGGGCTAGCGAATTCGAATGATTGGATTGTCGTGG | Forward primer to clone AN400_RS26690 with a C-termina $His_6$ tag in pBAD18 by NEBuilder HiFi DNA Assembly. Primer is used to linearise vector |
| | AGGATCCCCGGGTACCGAGCTTAGTGATGGTGATGGTG | Forward primer to clone AN400_RS26690 with a C-termina $His_6$ tag in pBAD18 by NEBuilder HiFi DNA Assembly. Primer is used to linearise vector |

* For the following plasmids the full length insert was synthetised by GenScript and the genes of interest were cloned in pBAD18 by NEBuilder HiFi DNA Assembly

| Templates for cell-free *in vitro* protein synthesis | Sequence of relevant primers (5'-3')[a] | Description |
|---|---|---|

| | | |
|---|---|---|
| ShdA II | GCGAATTAATACGACTCACTATAGGGCTTAAG<br>TATAAGGAGGAAAAAATATGCTGCAGGATCGC<br>TTTGCGGCCC | Forward primer to generate a DNA template for the *in vitro* synthesis of ShdA II (AN400_RS26690) from aminoacid 138 to 524 |
| | AAA CCC CTC CGT TTA GAG AGG GGT TAT<br>GCT AG TTA<br>CACGCCTGCTGCTCAGCTGGCATC | Reverse primer to generate a DNA template for the *in vitro* synthesis of ShdA II (AN400_RS26690) from aminoacid 138 to 524 |
| ShdA I | GCGAATTAATACGACTCACTATAGGGCTTAAG<br>TATAAGGAGGAAAAAAATGCTGCAGGATCGCT<br>TCGCGGCCCTG | Forward primer to generate a DNA template for the *in vitro* synthesis of ShdA I (DL351_RS13220) from aminoacid 140 to 526 |
| | AAACCCCTCCGTTTAGAGAGGGGTTATGCTAT<br>TACTGCGCCATTTCGCTGGACTC | Reverse primer to generate a DNA template for the *in vitro* synthesis of ShdA I (DL351_RS13220) from aminoacid 140 to 526 |
| ShdA III | GCGAATTAATACGACTCACTATAGGGCTTAAG<br>TATAAGGAGGAAAAAAATGCAGGACAAATTCG<br>CCAACCTGTC | Forward primer to generate a DNA template for the *in vitro* synthesis of ShdA III (EQ826_RS12590) from aminoacid 135 to 523 |
| | AAACCCCTCCGTTTAGAGAGGGGTTATGCTAG<br>TTATGCTTCCTCCTGCGCCTCGAAGGC | Reverse primer to generate a DNA template for the *in vitro* synthesis of ShdA III (EQ826_RS12590) from aminoacid 135 to 523 |
| ShdA IV | GCGAATTAATACGACTCACTATAGGGCTTAAG<br>TATAAGGAGGAAAAAAATGCAGGATAAGTTCG<br>CCAATCTGTCCAAG | Forward primer to generate a DNA template for the *in vitro* synthesis of ShdA IV (A9179_RS12860) from aminoacid 135 to 521 |
| | AAACCCCTCCGTTTAGAGAGGGGTTATGCTAG<br>TTACTGCGGCAATTCGCTGGGCTCATG | Reverse primer to generate a DNA template for the *in vitro* synthesis of ShdA IV (A9179_RS12860) from aminoacid 135 to 521 |

4. B. Ize, *et al.*, Remnant signal peptides on non-exported enzymes: implications for the evolution of prokaryotic respiratory chains. *Microbiology* **155**, 3992–4004.
